# Asgard archaea modulate potential methanogenesis substrates in wetland soil

**DOI:** 10.1101/2023.11.21.568159

**Authors:** Luis E. Valentin-Alvarado, Kathryn E. Appler, Valerie De Anda, Marie C. Schoelmerich, Jacob West-Roberts, Veronika Kivenson, Alexander Crits-Christoph, Lynn Ly, Rohan Sachdeva, David F. Savage, Brett J. Baker, Jillian F. Banfield

## Abstract

The roles of Asgard archaea in eukaryogenesis and marine biogeochemical cycles are well studied, yet their contributions in soil ecosystems are unknown. Of particular interest are Asgard archaeal contributions to methane cycling in wetland soils. To investigate this, we reconstructed two complete genomes for soil-associated Atabeyarchaeia, a new Asgard lineage, and the first complete genome of Freyarchaeia, and defined their metabolism *in situ*. Metatranscriptomics highlights high expression of [NiFe]-hydrogenases, pyruvate oxidation and carbon fixation via the Wood-Ljungdahl pathway genes. Also highly expressed are genes encoding enzymes for amino acid metabolism, anaerobic aldehyde oxidation, hydrogen peroxide detoxification and glycerol and carbohydrate breakdown to acetate and formate. Overall, soil-associated Asgard archaea are predicted to be non-methanogenic acetogens, likely impacting reservoirs of substrates for methane production in terrestrial ecosystems.

**One-Sentence Summary:** Complete genomes of Asgard archaea, coupled with metatranscriptomic data, indicate roles in production and consumption of carbon compounds that are known to serve as substrates for methane production in wetlands.

## Introduction

Wetland soils are hotspots for methane production by methanogenic archaea. The extent of methane production depends in part on the availability of substrates for methanogenesis (e.g., formate, formaldehyde, methanol, acetate, hydrogen), compounds that are both produced and consumed by co-existing microbial community members. Among the groups of organisms that coexist with methanogens are Asgard archaea, of recent interest from the perspective of eukaryogenesis (*1–4*). To date, numerous lineages of Asgard archaea have been reported from anaerobic, sedimentary freshwater, marine, and hydrothermal environments (*1–15*). Predictions primarily from draft metagenome-assembled genomes (MAGs) indicate metabolic diversity and flexibility that may enable them to occupy these diverse ecological niches. It appears that Asgard archaea are not capable of methane production since they lack the key canonical methyl-coenzyme M reductase (MCR). Although a few complete genomes for Asgard from hydrothermal and geothermal environments have been reported (*9*, *15–17*), most metabolic analyses of Asgard archaea are limited by reliance on partial genomes. To date, no Asgard genomes from non-estuarine wetland soils have been reported. Thus, nothing is known about the ways in which Asgard archaea directly (via methane production) or indirectly (via metabolic interactions) impact methane cycling in wetlands.

To investigate the roles of Asgard archaea in carbon cycling in wetland soil, we reconstructed two complete genomes for a newly defined group, here named Atabeyarchaeia, and one complete genome for a group named Freyarchaeia. Freyarchaeia MAGs were originally reconstructed from Guaymas Basin, located in the Gulf of California, México (*14*), and from Jinze Hot Spring (Yunnan, China) (*4*). Subsequently, another group used the original data to recover similar genomes and referred to them as Jordarchaeia (*18*). Here, we retain the original nomenclature. The genomes for soil Asgard archaea were initially reconstructed by manual curation of Illumina short read assemblies and then validated using both Nanopore and PacBio long reads. These fully curated genomes enabled us to perform comprehensive metabolic analyses, without the risks associated with reliance on draft genomes, and provided context for metatranscriptomic measurements of their *in situ* activity. Our integrated analysis of gene expression and metabolic predictions revealed roles for Atabeyarchaeia and Freyarchaeia in the production and consumption of carbon compounds that can serve as substrates for methanogenesis by coexisting methanogenic archaea.

## Results

### Complete genomes and phylogenetic placement of Asgard archaea from wetland soil

We analyzed Illumina metagenomic data from samples collected from 20 cm to 175 cm depth in the soil of a wetland located in Lake County, California, USA. We previously reported megaphages (*19*) and *Methanoperedens* archaea and their 1 Mb-scale “Borg” extrachromosomal elements from this site (*20*). From the metagenomic analyses conducted at this site, we determined that archaea account for >45% of the total community below a depth of 60 cm. Archaeal groups detected include members of the Asgardarchaeota, Bathyarchaeia, Methanosarcinia, Nitrososphaeria, Thermoplasmata, Micrarchaeia, Diapherotrites, Aenigmatarchaeia, Methanomicrobia, Aenigmarchaeia, Nanoarchaeia, Hadarchaeia and Methanomethylicia (**Fig. 1A-B**)

**Figure 1.**
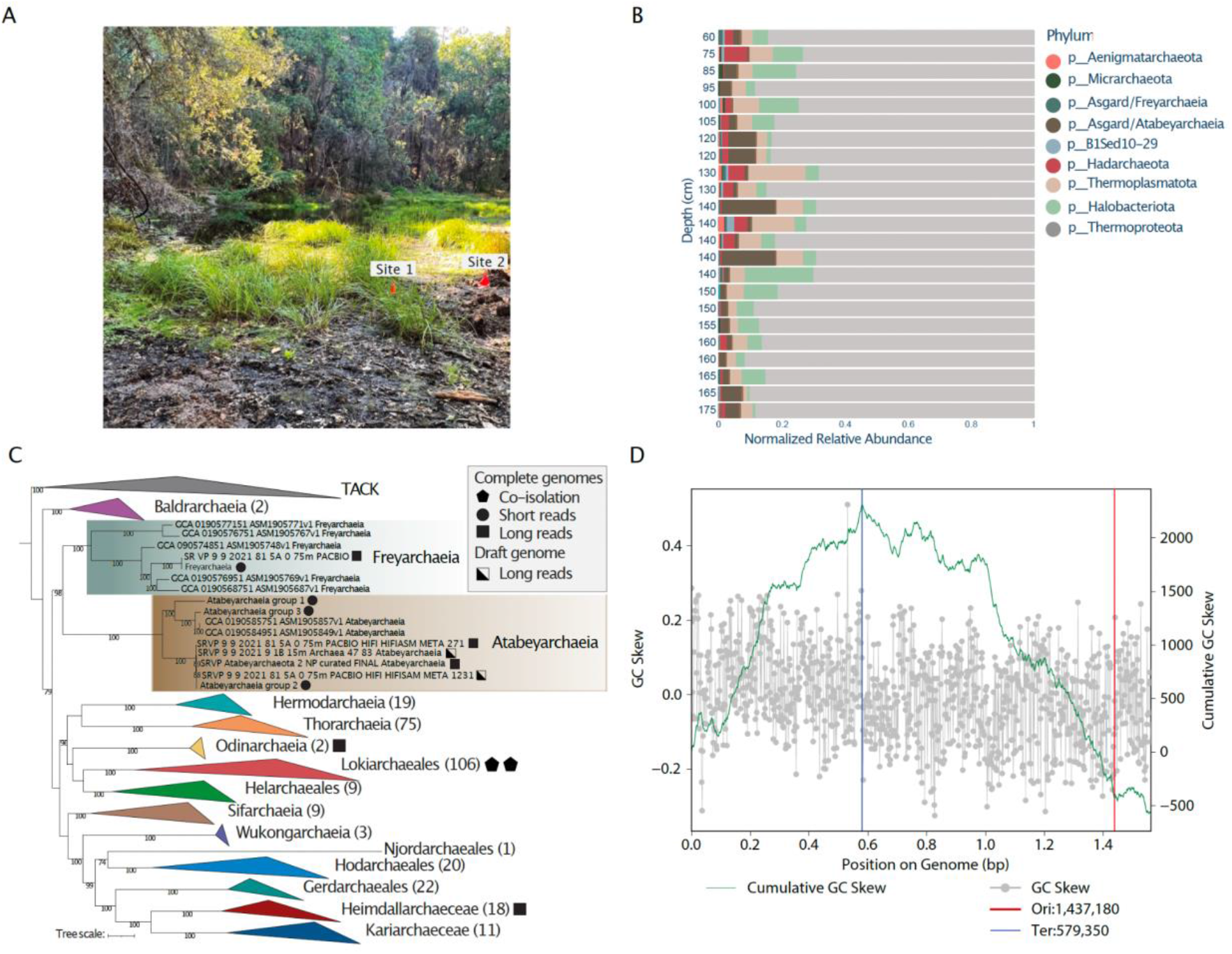
Archaea dominate deep regions of wetland soil and host novel Asgard archaea. **A.** Photograph of the vernal pool that was metagenomically sampled in this study, in Lake County, California, USA. **B.** Archaeal genomic abundance excluding bacterial genomes. **C.** Phylogenetic distribution of Asgard Archaea complete genomes. The maximum-likelihood phylogeny was generated with Iqtree v1.6.1, utilizing 47 concatenated archaeal Clusters of Orthologous Groups of proteins (arCOGs). The best-fit model was determined as LG+F+R10 based on the Bayesian Information Criterion. Non-parametric bootstrapping was conducted with 1,000 replicates for robustness. The filled-in square, circle, and triangle indicate closed complete genomes from short reads, published complete genomes from long reads, and genomes from co-isolated cultured representatives, respectively. The pentagon highlights the long read draft genomes from this site (PacBio or Nanopore). **D.** Bidirectional replication indication in Atabeyarchaeia complete genomes. The GC skew is shown as a grey plot overlaying the cumulative GC skew, presented as a green line. The blue lines mark the predicted replication terminus.

From 60 cm, 80 cm, and 100 cm deep wetland soil, we recovered four draft Asgard genomes, three of which were manually curated to completion using methods described previously (*21*). Taxonomic classification using the Silva DB placed the 3,576,204 bp genome as Freyachaeia. 16S rRNA gene sequence analysis showed the two other complete genomes were distinct from Freyarchaeia (16S rRNA genes are <75% identical), thus representing organisms from a separate, new lineage. These genomes are 2,808,651 and 2,756,679 bp in length (**table S1**) with an average amino acid identity (AAI) of ~70% (**table S2**).

Phylogenetic analyses using several sets of marker genes (**“see materials and methods”**) placed our two novel complete genomes in a monophyletic group within the Asgard clade as a sister group to Freyarchaeaia (**Fig. 1C**). **We** performed phylogenetic analyses using concatenated marker sets of 47 arCOG and 15 ribosomal protein (RP15) gene cluster (**fig. S1)**, as well as 16S rRNA (**fig. S2)**. The new genomes share only 40-45% AAI when compared to other Asgard genomes, consistent with their assignment to a new phylum. Although our analyses provide evidence for distinction at the phylum level, we chose to adhere to the Genome Taxonomy Database (GTDB) for standardized microbial genome nomenclature (**table S3**). Here, we propose the name *Candidatus* “Atabeyarchaeia’’ for this new group, where ‘Atabey’ is a goddess in of Taíno Puerto Rican mythology. Atabeyarchaeia is represented by the complete Atabeyarchaeia group 1 (Atabeya-1) and group 2 (Atabeya-2) genomes. Included in this group are 2 MAGs from a highly fragmented, partial Asgard Lake Cootharaba Group (ALCG) draft genome (*12*). The cumulative GC skew of the Freyarchaeia and Atabeyarchaeia genomes is consistent with bidirectional replication. This style of replication is typical of bacterial genomes but has not been widely reported in Archaea, and has never been described in the Asgard group (**Fig. 1D and fig. S3**).

Unexpectedly, we found that 92% to 95% of tRNA genes from all three genomes contain at least one intron. This contrasts with the general estimate that 15% of archaeal tRNA harbor introns (*22*), and with Thermoproteales (another order of archaea), where 70% of the tRNAs contain introns (*23*). In total, there are 228 tRNA introns across the three new Asgard genomes (**table S4**). Unlike most archaeal tRNA introns that occur in the anticodon loop at position 37 / 38 (*24*, *25*), Atabeyarchaeia and Freyarchaeia introns often occur at non-canonical positions, and over half of their tRNA genes have multiple introns (**table S4**).

Subsequently, we acquired and independently assembled Oxford Nanopore and PacBio long-reads from a subset of the samples to generate three circularized genomes that validate the overall topology of all three curated Illumina read-based genomes (**fig. S4, table S1**). These complete genomes allowed us to genomically describe two Atabeya-2 strain variants from 100 cm and 175 cm depth soil. In addition, we used Illumina reads to curate a draft Nanopore genome for another Atabeyarchaeia species, Atabeya-3, from 75 cm and 175 cm depth soil (**fig. S5**). The Atabeyarchaeia-3 genome is most closely related to the Asgard Lake Cootharaba Group (ALCG) fragments (*18*). To further solidify the phylogenetic position of Atabeyarchaiea, we included the Atabeyarchaeia-3 genome and another draft genome (Atabeyarchaeia-4) from Illumina reads in the phylogenetic analysis.

Using the Asgard clusters of orthologous genes (AsCOGS) database and functional classification, we identified eukaryotic signature proteins (ESPs) in the complete and public genomes of Atabeyarchaeia and Freyarchaeia (*2*, *3*). Atabeyarchaeia and Freyarchaeia genomes had the highest percentage of hits for ‘Intracellular trafficking, secretion, and vesicular transport’ (U) among the AsCOG functional classes, accounting for 84.3% of the hits to the database. Within this class, we identified key protein domains such as Adaptin, ESCRT-I-III complexes, Gelsolin family protein, Longin domain, Rab-like GTPase, Ras family GTPase, and Roadblock/LC7 domain **(table S5, fig. S6**). The ‘Post Translational modification, protein turnover, and chaperones’ category (O) followed with a count of 101 (15.8%), highlighting domains like Ubiquitin, Jab1/MPN domain-containing protein, and the RING finger domain. The presence of ESPs in the newly described Atabeyarchaeia lineage and their presence in Freyarchaeia aligns with previous findings for Asgardarchaeota (*1*, *3*, *4*).

### Expression of energy conservation pathways constrain key metabolisms in situ

We analyzed the metabolic potential of the three complete genomes and investigated their activity *in situ* through metatranscriptomics of soil samples (**“see materials and methods”**, **Fig. 2**, **table S6, table S7**). The metatranscriptomic data indicate high expression of genes involved in key energy conservation pathways (**Fig. 3A**). Most highly transcribed genes are soluble heterodisulfide reductase (HdrABC), [NiFe] hydrogenases (groups 3 and 4), ATP synthase, numerous aldehyde ferredoxin oxidoreductase genes, genes for phosphoenolpyruvate (PEP) and pyruvate metabolism, and carbon monoxide dehydrogenase/acetyl CoA synthetase (CODH/ACS). Notably, the Hdr, the group 3 and group 4 hydrogenase (including up to eight NADH-quinone oxidoreductase subunits, e.g., Nuo-like) as well as the ATP synthase are co-encoded in a syntenic block in all of the genomes (**Fig. 4A**). Phylogenetic analysis of the large subunit of group 4 [NiFe]-hydrogenases suggests they are closely related to those of Odinarchaeia, Heimdallarchaia, and Hermodarchaeia (**Fig. 4B, table S8**). However, the exact function of this unclassified Asgard group has not been validated biochemically (*26*). One clue relies on the identification of eight genes homologous to the hydrophobic subunits of complex I NuoL, M, and N (*E. coli* nomenclature) and Mrp-type Na+/H+ antiporters. Thus, these Asgard archaea may mediate Na+/H+ translocation coupled to energy generation via ATP synthase (*27–29*).

**Figure 2.**
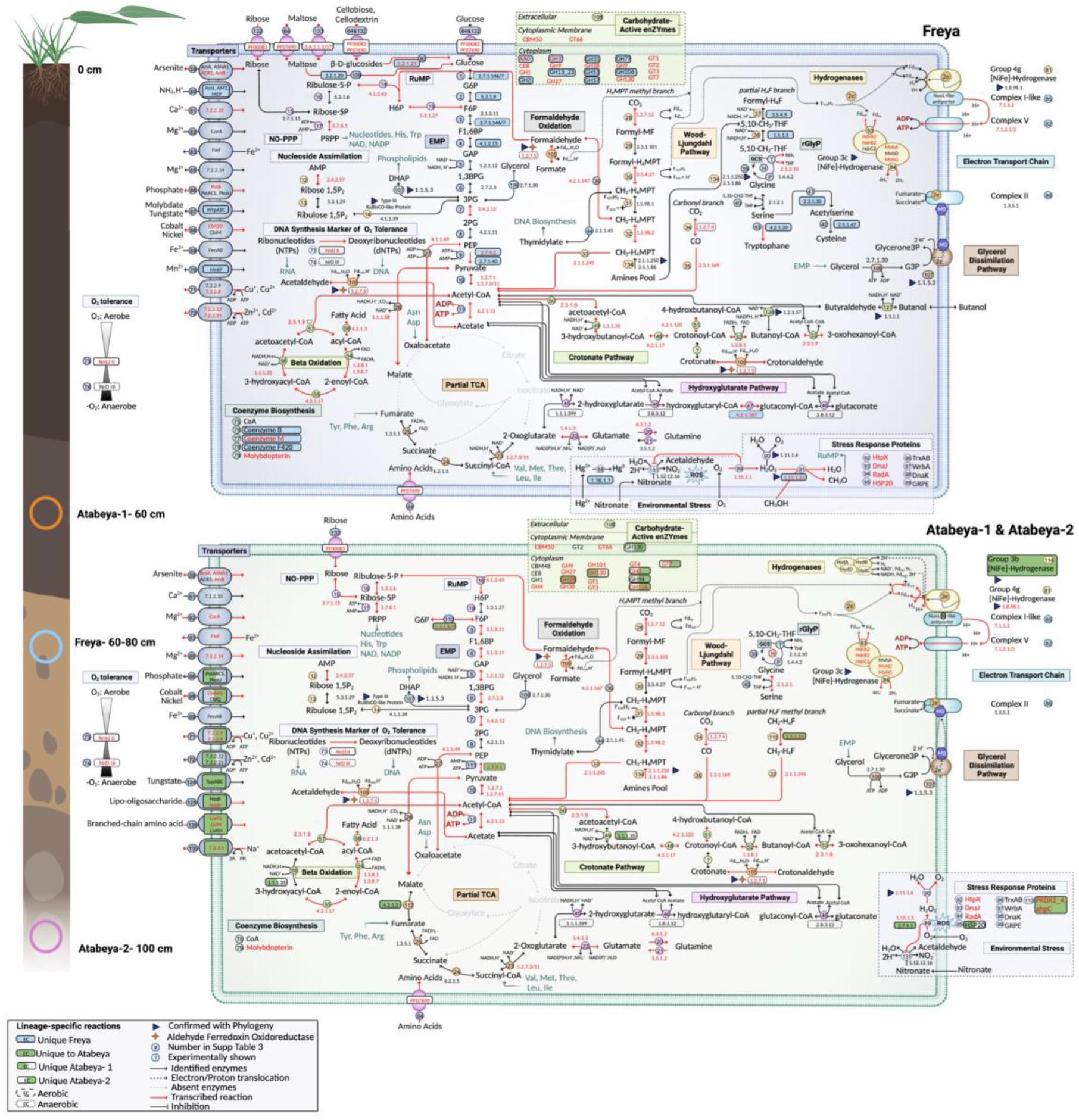
Metabolic capacities of terrestrial Atabeyarchaeia and Freyarchaeia for overall implications for biogeochemical cycling in wetlands. Inference of the pathways from the complete genomes is based on the comparison of predicted proteins with a variety of functional databases (**“see materials and methods”**). The extraction depth location within the cores is shown on the left. All reactions are numbers and correspond to **table S7**. EC/TCDB numbers shaded fully or partially in blue or green are unique to the lineages and complete genomes, whereas the dashed boxes distinguish oxygen-sensitive enzymes. The multi-functional aldehyde ferredoxin oxidoreductase is shown with a star. Proteins marked with a triangle have generated phylogenies to determine their evolutionary histories and substrate specificity. Reactions with mapped transcripts are denoted with red text and arrows. Created using BioRender.com.

**Figure 3.**
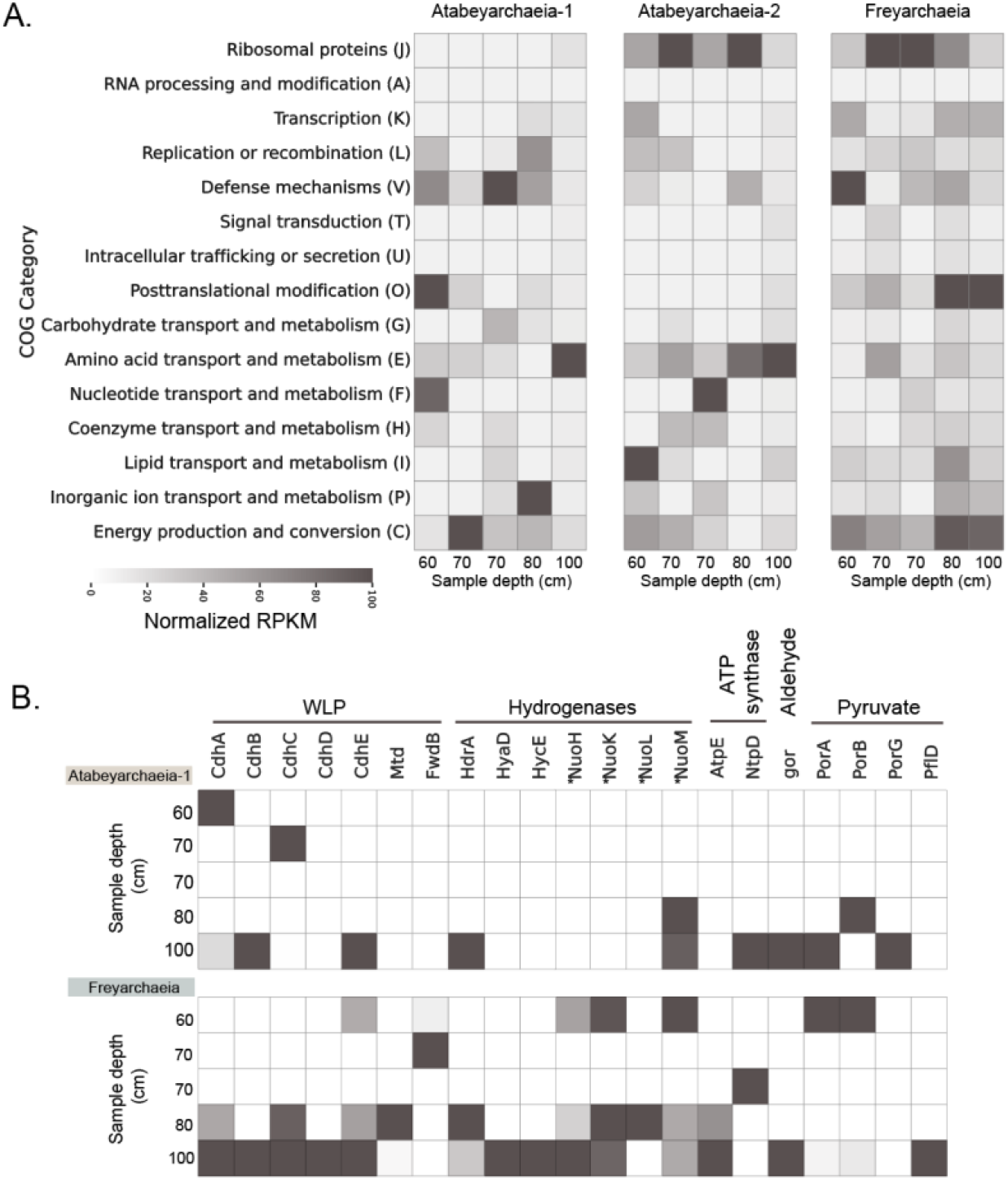
Metatranscriptomic profiling of soil-associated Asgard archaeal genomes. **A.** Heatmap visualization of normalized Reads Per Kilobase per Million mapped reads (RPKM) values for ORFs with high sequence similarity (≥95%) to the genomes of Atabeyarchaeia-1, Atabeyarchaeia-2, and Freyarchaeia, across various soil depths. A total of 2,191 open reading frames (ORFs) were categorized using the Clusters of Orthologous Groups (COG) database, with Atabeya-1, Atabeya-2, and Freya expressing 465, 804, and 922 unique ORFs, respectively. The ORFs were annotated and assigned to 15 COG categories, indicating the functional potential of each archaeal genome in situ. Columns represent metatranscriptomes from different soil depths, highlighting the spatial variability in the expression of key metabolic and cellular processes. **B.** Expanded heatmap of Atabeyarchaeia-1 and Freyarchaeia expressed genes under the category C: Energy production and conversion. Key genes of the WLP (CODH/ACS, carbon monoxide dehydrogenase/acetyl-CoA synthase; *fwdB*, formate dehydrogenase; mtd, 5,10-methylene-H4-methanopterin dehydrogenase), hydrogenases and associated genes (HdrA, heterodisulfide reductase and group NiFe-hydrogenase; Mvh, methyl viologen reducing hydrogenase); HyaD (NiFe-hydrogenase_maturation_factor); HycE and Nuo like subunits, (group 4 NiFe-hydrogenase), ATP synthase (AtpE, V/A-type H+/Na+-transporting ATPase subunit_K; NtpD, V/A-type H+/Na+ transporting ATPase subunit D) and aldehyde metabolism (gor, Aldehyde:ferredoxin oxidoreductases), pyruvate oxidation (porABCD, 2-pyruvate:ferredoxin oxidoreductase; pflD, pyruvate-formate lyase).

**Figure 4.**
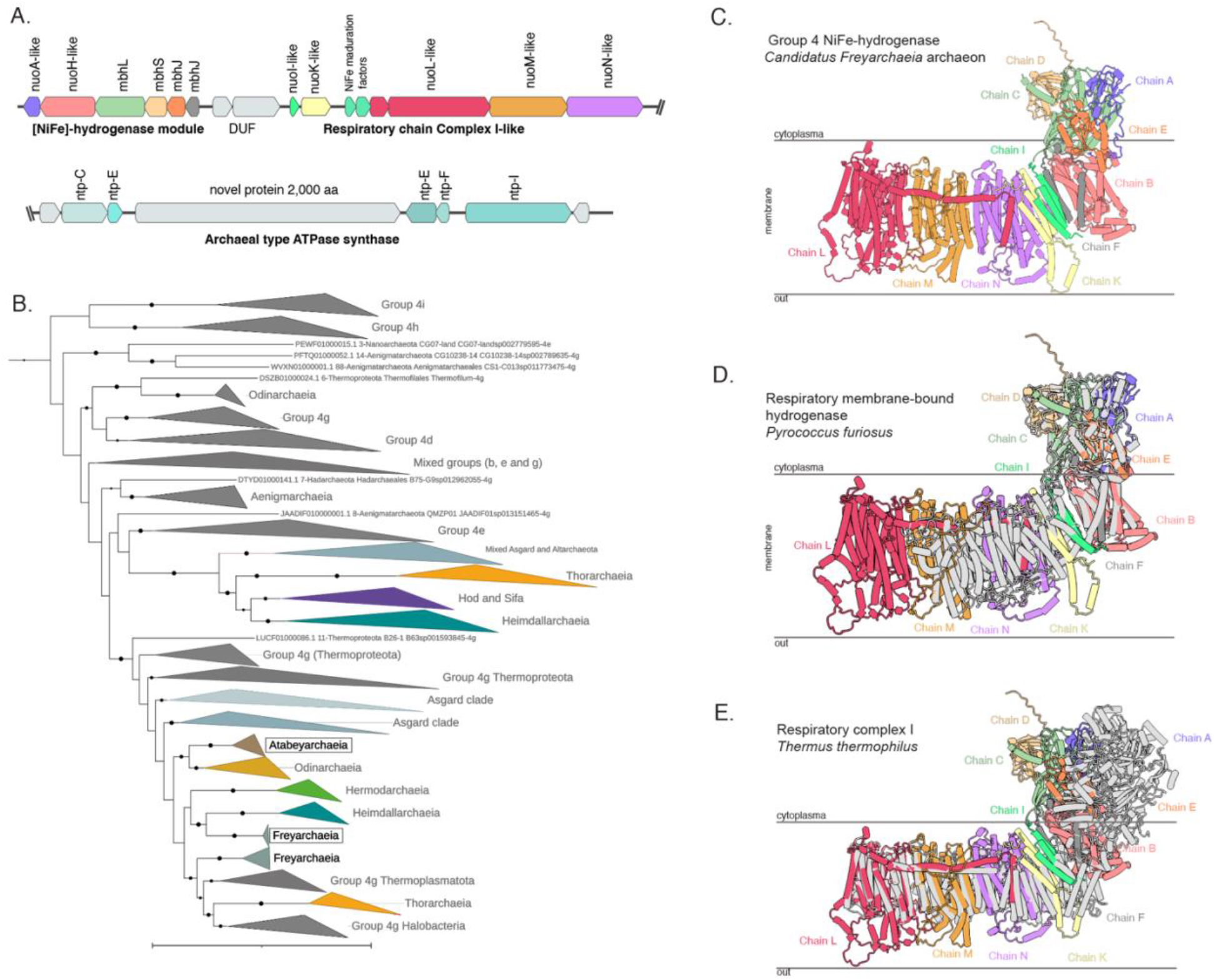
Phylogeny, genetic organization and structure of the novel group 4 energy-conservation complex I-like NiFe-hydrogenase from Asgard archaea. **A.** Genetic organization of the group 4 [NiFe]-hydrogenase module, the proton-translocating membrane module, and ATP synthase from the Freyarchaeia genome. **B.** Maximum likelihood phylogeny of group 4 [NiFe]-hydrogenase large subunit from Asgard archaea and reference sequences. The bolded taxonomic groups highlight the clades with genomes from this study used for modeling. **C.** AlphaFold models of [NiFe]-hydrogenase module and the proton-translocating membrane module where each candidate subunit is represented by a different color based on the best subunit matched. **D.** AlphaFold model of Freyarchaeia hydrogenase complex colored by chains, aligned with cryoEM structure of a respiratory membrane-bound hydrogenase (MBH) from *Pyrococcus furiosus* (*27*) (PDB ID: 5L8X). E. AlphaFold model of Freyarchaeia hydrogenase complex colored by chains, aligned with Crystal structure of respiratory complex I from *Thermus thermophilus*(*31*) (PDB: 4HEA).

We employed AlphaFold2 to model the hydrogenase and associated complex I-like modules. Overall, the predicted structure has a cytosolic and membrane-associated portion (**Fig. 4C**). The cytosolic portion aligned with the respiratory membrane-bound hydrogenase (MBH) from *Pyrococcus furiosus* (*27*) with high confidence (**Fig. 4D**). When superimposed, the calculated structures of the membrane-associated hydrophobic L, M, K, and S chains aligned to bacterial complex I. In the canonical complex I (*30*, *31*), Chain L, Nqo12, as well as M, N, and K translocate proteins (*31*, *32*), a process that is facilitated by an arm, helix HL that is part of chain L. This helix HL is also present in the L-like subunit of the Asgard complexes (**Fig. 4E**). The helix HL, and the antiporter subunits located between chain L and the subunit that connects to the cytosolic hydrogenase portion, are absent in all characterized respiratory membrane-bound hydrogenases (**Fig. 4E, fig. S7**).

The Group 3c cofactor-coupled bidirectional [NiFe] hydrogenase (**fig. S8A**) in combination with HdrABC suggests the capability to bifurcate electrons from H_2_ to ferredoxin and an unidentified heterodisulfide compound. This capacity has been observed in methanogenic archaea via the MvhADG–HdrABC system (*33*, *34*). Atabeyarchaeia genomes encode two independent gene clusters of the Group 3b NADP-coupled [NiFe] hydrogenases (**fig. S8B**). Their presence suggests the capacity to maintain redox equilibrium and, potentially, grow lithoautotrophically by using H_2_ as an electron donor, as suggested for other Asgardarchaeota members (*10*, *14*, *26*).

Atabeyarchaeia and Freyarchaeia encode both the tetrahydromethanopterin (H_4_MPT) methyl branch and the carbonyl branch of the Wood–Ljungdahl pathway (WLP) (**fig. S9**). This reversible pathway can be used to reduce CO_2_ to acetyl coenzyme A (acetyl CoA), which can be further converted to acetate. This last conversion can lead to energy conservation in both Asgard lineages via substrate-level phosphorylation when mediated by acetate-CoA ligase (see below). We confirmed the expression of almost all of the genes of the methyl and carbonyl branches, including the acetate-CoA ligase, in all complete genomes. When H_2_ is present in the ecosystem, these archaea could use the WLP for the reduction of CO_2_ or formate and thereby conserving energy. Alternatively, they could use the WLP in reverse to oxidize acetate. In both scenarios, the expression of energy-converting hydrogenases and the ATP synthases suggest a potential role in energy conservation. This involvement may include coupling exergonic electron transfer to establish an ion gradient that fuels the ATP synthase for ATP generation. The metabolic inferences along with the transcriptional data including the expression of *por* genes in all three Asgard genomes, indicates a reliance on an archaeal version of the WLP to perform acetogenesis (*34*, *35*). This acetogenic lifestyle appears to to involve energy conservation through a hydrogenase-dependent chemiosmotic mechanism similar to that observed in some acetogenic bacteria (*36*).

### Potential for non-methanogenic methylotrophic life-style and carboxydotrophy

Despite the absence of the MCR complex, Freyarchaeia genomes have all the necessary genes to synthesize coenzyme-M from sulfopyruvate via the ComABC pathway similar to methanogens (*37*, *38*). Most methanogens conserve energy via the Na+-translocating MtrA-H complex, which is encoded by an eight-gene cluster (*39*). Although Atabeyarchaeia and Freyarchaeia do not have the genes for the full complex, Atabeyarchaeia-1 has two copies of the CH_3_-H_4_MPT-dependent methyltransferase subunit A-like (MtrA) and both Freyarchaeia and Atabeyarchaeia also encode the CH_3_-H_4_MPT-dependent methyltransferase subunit H (MtrH), along with a phylogenetically distinct fused polypeptide of MtrA-like and MtrH (**Figure 5A**). Under the conditions prevalent at the time of sampling, the *mtr* genes were only weakly expressed (**table S6, S8**). While the biochemical activity of these divergent non-methanogen-associated MtrA-like and MtrH-like enzymes remain unclear, our phylogenetic analyses suggest they are phylogenetically related to methanogenic MtrA, MtrH, and MtrAH sequences. This suggests their potential role in converting CH_3_-H_4_MPT to H_4_MPT, transferring a methyl group to an acceptor –possibly coenzyme-M, which can be produced by Freyarchaeia–. As they lack the MCR complex, the subsequent fate of the methyl group remains uncertain.

**Figure 5.**
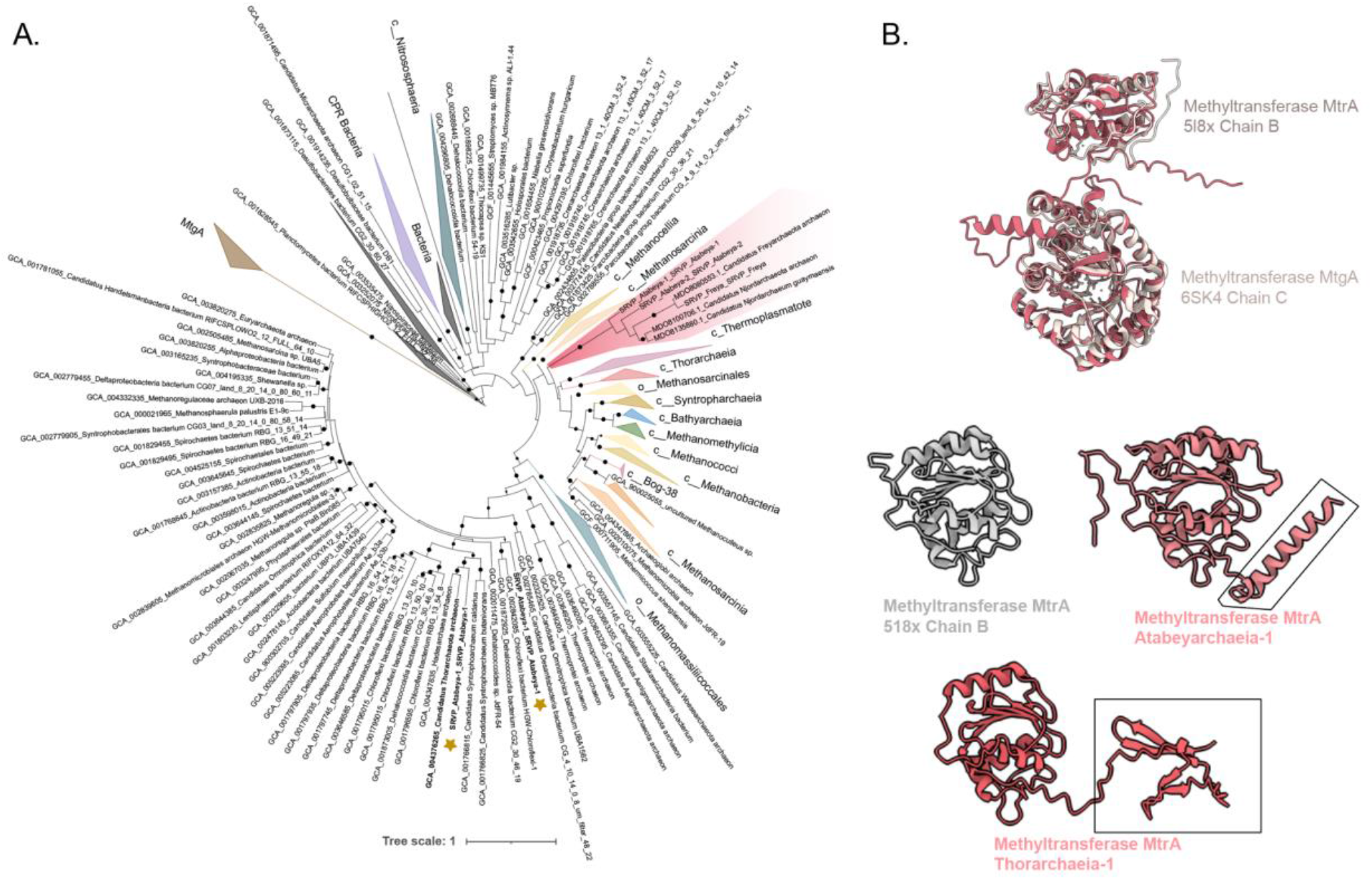
Non-methanogenic MtrA, MtrH and MtrAH fusion methyltransferases. **A.** Maximum likelihood phylogeny of MtrA and the MtrAH fusion, with reference to Tetrahydromethanopterin S-methyltransferase subunit A (MtrA) with the closest corresponding domains being MtrA from the characterized Tetrahydromethanopterin S-methyltransferase subunit A (MtrA) protein (PDB ID: 5L8X) (*67*). The coral colored clade is the novel fusion present in Atabeyarchaeia, Freyarcheia and other Asgardarchaeota members. **B.** AlphaFold models of Atabeyarchaeia-1 MtrAH (fusion) in coral aligned with the grey corresponding domains of the characterized protein Tetrahydromethanopterin S-methyltransferase subunit A (MtrA) (PDB ID: 5L8X)(*68*) and Methyltransferase (MtgA) from *Desulfitobacterium hafniense* in complex with methyl-tetrahydrofolate (PDB ID: 6SK4) at the N terminus. We also modeled the putative MtrA present in Atabeyarchaeia-1 with the closest corresponding domains being MtrA from the characterized Tetrahydromethanopterin S-methyltransferase subunit A (MtrA) protein (PDB ID: 5L8X).

**Figure 6.**
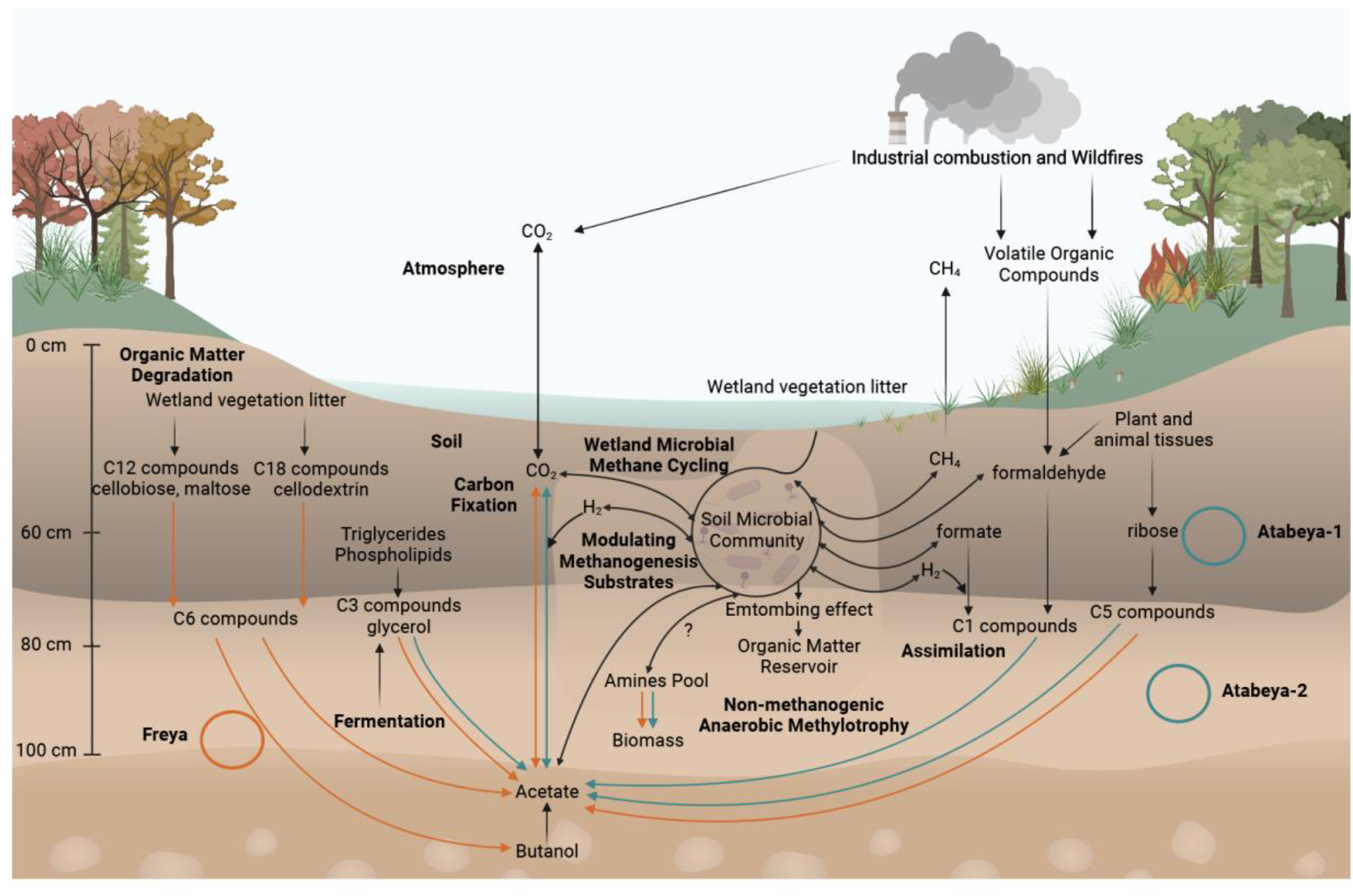
Overview of the wetland soil dynamics and biogeochemical cycling in Atabeyarchaeia and Freyarchaeia. Complete genomes for Atabeyarchaeia and Freyarchaeia are shown with green and orange circles, respectively. 2 Atabeyarchaeia genomes (Atabeya-1 and Atabeya-2) and 1 Freyarchaeia (Freya) genome were isolated and carefully curated and closed from wetland soil between 60-100 cm. These anaerobic lineages were shown in this study to encode the Wood-Ljungdahl Pathway for CO_2_ fixation (e.g. methylated compounds such as quaternary amines) and EMP Pathway, components of chemolithotrophy and heterotrophy, producing acetate shown in arrows (green and orange), corresponding to the genome colors. Additionally, these lineages are involved in modulating methanogenesis substrates in these wetland soils. Detailed description of the specific pathways is found in main text, Fig. 2, and supplementary materials. Created using BioRender.com.

Although Atabeyarchaeia and Freyarchaeia genomes do not encode MrtE, we identified genes associated with methyltransferase systems encoded in close proximity to the MtrH gene. Specifically, the genomes encode trimethylamine methyltransferase (MttB-like, COG5598 superfamily), undefined corrinoid protein (MtbC-like), and putative glycine cleavage system H (gcvH) (**table S9**). Both Atabeyarchaeia and Freyarchaeia genomes encode trimethylamine methyltransferase MttB (COG5598). Phylogenetic analysis suggests that MttB (**fig. S10**) and MtbC **(fig. S11)** belong to a previously uncharacterized group of methyltransferases, similar to those found in Njordarchaeales, Helarchaeales, Odinarchaeia and TACK members, including Brockarchaeia and Thermoproteota. In methanogens that encode *mttB*, this gene has an amber codon encoding the amino acid pyrrolysine in the active site (*40*, *41*). The archaea from this study do not encode pyrrolysine, suggesting Freyarchaeia and Atabeyarchaeia encode a non-pyrrolysine MttB homolog, likely a quaternary amine (QA) dependent methyltransferase (*42*). Only a fraction of QA methyltransferase substrates have been identified, and these include glycine betaine, proline betaine, carnitine, and butyrobetaine (*42–45*). The methyl group from the QA may be transferred to THF or H_4_MPT branches of the WLP, akin to the mechanisms described in archaea with the capacity for non-methanogenic anaerobic methylotrophy, including Freyarchaeia (Jordarchaeia), Sifarchaeia, Brockarchaeia, and Culexarchaeia (*11*, *12*, *46*, *47*). Consumption of QA compounds may reduce the pool of potential substrates for methanogenic methane production.

We identified genes in the Freyarchaeia genome that potentially encode an aerobic carbon-monoxide dehydrogenase complex (CoxLMS) and associated cofactors. Phylogenetic analysis places the putative CoxL in a monophyletic group with other archaea including Thermoplasmatales, Marsarchaeota, and Culexarchaeia (**fig. S12**). The gene cassette arrangement suggests these archaea may possess the ability to use carbon monoxide as a growth substrate (carboxydotrophy). However, analysis of the protein sequence reveals that the putative large-subunit aerobic CO dehydrogenases (CoxL) are missing the characteristic VAYRCSFR motif, which is critical for CO binding in the form I Cox proteins (*48*, *49*). Nevertheless, the modeled protein structure, along with the operon organization of the *cox* genes, points to a novel type of Cox system in archaea (**fig. S13**). Alternatively, it is possible that this complex enables the utilization of alternative substrates, such as aldehydes or purines, as a member of the aldehyde oxidase superfamily (*47*, *49*, *50*).

### Carbon compound metabolic pathways

There are indications that Freyarchaeia and Atabeyarchaeia display distinct metabolic preferences for various soil carbon compounds (**Fig. 2; fig. S13**). Freyarchaeia exhibit a genetic repertoire to break down various extracellular lignin-derived compounds including 5-carboxyvanillate. Other substrates that we predict can be metabolized by Freyarchaeia carbohydrate-active enzymes include hemicellulose (C5), cellobiose, maltose, and cellulose (C12). We predict that cellodextrin (C18) compounds can be converted to glucose via beta-glucosidase (BglBX). The findings implicate Freyarchaeia in the metabolism of plant-derived soil carbon compounds. Glucose, resulting from the degradation of complex carbohydrates, as well as ribulose and other carbon substrates, likely enters the modified Embden-Meyerhof-Parnas (EMP) pathway, yet the genes of this EMP pathway genes are only weakly expressed (**fig. S14, table S6, S8)**. Additionally, Freyarchaeia encode and express an array of genes for the uptake of carbohydrates including major facilitator superfamily sugar transporters and ABC-sugar transporters suggesting an active role in efficiently assimilating diverse carbon substrates from soil environments Atabeyarchaeia also harbor genes of the EMP glycolytic pathway, producing ATP through the conversion of acetyl-CoA to acetate (**fig. S13**). Unlike Freyarchaeia which likely feed glucose into the EMP pathway, the entry point for Atabeyarchaeia to the EMP pathway appears to be fructose 6-phosphate (F6P). This is relatively uncommon for Asgard archaea but is reminiscent of the pathway in Helarchaeales (*7*), an order of Lokiarchaeia. We identified Atabeyarchaeia transcripts for all but one of the genes for the steps from G6P to acetate (table S8).

Atabeyarchaeia and Freyarchaeia utilize different enzymes to produce pyruvate. Atabeyarchaeia encode the oxygen-sensitive reversible enzyme, pyruvate:phosphate dikinase (PpdK); whereas Freyarchaeia encodes unidirectional pyruvate water dikinase/phosphoenolpyruvate synthase (PpS) and pyruvate kinase (Pk), producing phosphoenolpyruvate and pyruvate (*51*), respectively. Pyruvate generated via EMP pathway can be then converted to acetyl-CoA by pyruvate:ferredoxin oxidoreductase (PorABCDG) complex using a low-potential electron carrier such as a ferredoxin as the electron donor. Alternatively, acetyl-CoA can also be generated via pyruvate formate-lyase (pflD) generating formate as a byproduct. The final step involves the conversion of acetyl-CoA to acetate via acetate-CoA ligase (ADP-forming) producing ATP via substrate level phosphorylation-a crucial energy conserving step during fermentation of carbon compounds in both lineages.

Lacking the ability to phosphorylate C6 carbon sources, Atabeyarchaeia converts ribulose-5-phosphate (C5) and fixes formaldehyde (C1) into hexulose-6-phosphate (H6P) via the ribulose monophosphate (RuMP) and non-oxidative pentose phosphate (NO-PPP) pathways (**Fig.2, fig. S14**). The Atabeyarchaeia RuMP pathway bifunctional enzymes (HPS-PH and Fae-HPS) are common in archaea and similar to methylotrophic bacterial homologs (*52*). The RuMP pathway in these Asgard archaea can modulate the formaldehyde availability, a byproduct of methanol oxidation, microbial organic matter decomposition, and combustion. High expression of aldehyde-ferredoxin oxidoreductases (AOR) genes suggest another mechanism for the interconversion of organic acids to aldehydes. For example, aldehyde detoxification (e.g., formaldehyde to formate) and source of acetate from acetaldehyde Atabeyarchaeia-1, Atabeyarchaeia-2, and Freyarchaeia encode multiple AOR gene copies 5, 6, and 8 respectively. Phylogenetic analyses (**fig. S15**) suggest that both Asgard lineages encode AOR genes related to the FOR family that oxidize C1-C3 aldehydes or aliphatic and aromatic aldehydes (e.g. formaldehyde or glyceraldehyde) (*53–55*). Furthermore, Freyarchaeia also encodes a tungsten-based AOR-type enzyme (XOR family) found in cellulolytic anaerobes with undefined substrate specificity (*56*) (**fig. S15**). Of the classified AORs, only one gene is expressed in Atabeyarchaeia-2 (**Figure 3**). Yet, some of the unclassified AOR genes are among the most highly expressed genes in the Atabeyarchaeia genomes. Despite the lack of biochemical characterization for most AOR families, these observations suggest a key role of multiple aldehydes in the generation of reducing power in the form of reduced ferredoxin.

Similar to other Asgard archaea (*7*, *10*, *26*), Atabeyarchaeia and Freyarchaeia encode genes for the large subunit of type IV and methanogenic type III Ribulose 1,5-bisphosphate carboxylase (RbcL) (**fig. S16**) a key enzyme in the partial nucleotide salvage pathway. This pathway facilitates the conversion of adenosine monophosphate (AMP) to 3-phosphoglycerate (3-PG), potentially leading to further metabolism into acetyl-CoA (*57*).

Anaerobic glycerol (C3) metabolism by Atabeyarchaeia and Freyarchaeia is predicted based on the presence of glycerol kinase (GlpK), which forms glycerol-3-phosphate (3PG) from glycerol. 3PG (along with F6P) can be broken down via the EMP pathway or 3PG can be converted to dihydroxyacetone phosphate (DHAP) via GlpABC. DHAP can also serve as a precursor for sn-glycerol-1-phosphate (G1P), the backbone of archaeal phospholipids. Freyarchaeia have an extra GlpABC operon, the GlpA subunit of which clusters phylogenetically with GlpA of Halobacteriales, the only known archaeal group capable of glycerol assimilation (**fig. S17**).

All three genomes have a partial TCA cycle similar to other anaerobic archaeal groups such as methanogens (*58*). They encode succinate dehydrogenase, succinyl-CoA synthetase, 2-oxoglutarate ferredoxin reductase that are important intermediates for amino acid degradation (e.g., glutamate). Only Atabeyarchaeia can convert fumarate to malate via fumarate hydratase. The only portion of TCA cycle transcribed in any genome is 2-oxoglutarate/2-oxoacid ferredoxin oxidoreductase, which can produce reducing power in the form of NADH.

A clue suggesting that amino acids are an important resource for Atabeyarchaeia and Freyarchaeia is the high expression of genes for protein and peptide breakdown (**Figure 2**). All three organisms are predicted to have the capacity to break down fatty acids via beta oxidation including crotonate (short-chain fatty acid) via the poorly described crotonate pathway. Furthermore they encode some enzymes involved in fermenting amino acids to H+, ammonium, acetate, and NAD(P)H via the hydroxyglutarate pathway (**table S7**). The genomes also encode amino acid transporters and these are also highly transcribed in both archaeal groups. The ability to anaerobically degrade amino acids is consistent with predictions of the metabolism of the last Asgard common ancestor (*4*, *9*).

Additionally, Freyarchaeia and Atabeyarchaeia can reverse the step in the formyl branch of the WLP that transforms glycine into methylenetetrahydrofolate (methylene-THF). Methylene-THF may then be converted to methyl-THF and then to formyl-THF, producing reducing power **(Figure 2**). Ultimately, the methyl group may be used to form acetate via the WLP. Interestingly Atabeyarchaeia-2 and Freyarchaeia expressed methylenetetrahydrofolate reductase (MTHFR) that is homologous to the enzyme used in the bacterial WLP and also plays a role in folate biosynthesis.

### Environmental protection and adaptations

We predict that Atabeyarchaeia and Freyarchaeia are anaerobes expressing genes that encode oxygen-sensitive enzymes and proteins that protect against oxidative and other environmental stressors. Interestingly, all three organisms encode an ancestral version of clade I catalases (KatE) (**fig. S18**), Fe-Mn superoxide dismutase (SOD2), and unique to Freyarchaeia, a catalase-peroxidase (**fig. S19**) for protection against reactive oxygen species (ROS) (**fig. S20**). Previous analyses have described these expressed enzymes in acetogenic and sulfate-reducing bacteria and methanogenic archaea, but to our knowledge, not in Asgard archaea, indicating a potential adaptation to soil environments(*59*). We also identified transcription for other environmental and stress responses, including transporters (e.g., nickel, arsenite, magnesium, iron, and copper), and heat shock proteins.

We infer that Atabeyarchaeia and Freyarchaeia use selenocysteine (Sec), the 21st amino acid, due to the presence of the Sec-specific elongation factor and Sec tRNA in their genomes. Additional Sec components, including phosphoseryl-tRNA kinase (Pstk), Sec synthase (SecS), selenophosphate synthetase (SPS) genes, and multiple Eukaryotic-like Sec Insertion Sequences are also present (**table S10**). Phylogenetic analysis shows that the Sec elongation factor sequences from Atabeyarchaeia and Freyarchaeia are closely related to other Asgard members and Eukaryotes (**fig. S21**). We identified multiple selenoproteins encoded within each genome, including CoB-CoM heterodisulfide reductase iron-sulfur subunit (HdrA), peroxiredoxin family protein (Prx-like), selenophosphate synthetase (SPS), and the small subunit (~50 aa) of NiFeSec (VhuU). In VhuU, Sec plays a crucial role in mitigating oxidative stress (55). Sec can also enhance the catalytic efficiency of redox proteins (56, 58), and the identified selenoproteins have the characteristic CXXU or UXXC sequence (**table S11**) observed in redox-active motifs (57).

## Discussion

Here, we reconstructed and validated three complete Asgard archaeal genomes from wetland soils in which these archaea comprise less than 1% of complex microbial communities. We used these genomes to define their chromosome lengths, structure and replication modes. It is relatively common for authors to report circularized genomes as complete, but this may be erroneous due to the prominence of local assembly errors, chimeras, scaffolding gaps and other issues in de novo metagenome assemblies (*21*, *60*). Our genomes were thoroughly inspected, corrected and vetted after circularization, steps previously described to complete genomes from metagenomes (*61*). These complete genomes are one of the first manual curations of short-read metagenomic data verified entirely with long-read analysis (Oxford Nanopore and/or PacBio) and the first complete short-read environmental Asgard genomes. Two of these genomes are from Atabeyarchaeia, a previously undescribed Asgard group and the first complete genome for Freyarchaeia. We predict bidirectional replication in Freyarchaeia and Atabeyarchaeia, suggesting bidirectional replication could have been present in the last common ancestor of eukaryotes and archaea, potentially playing a role in the emergence of the complex cellular organization characteristic of eukaryotes.

Overall, prior studies predict that Asgard archaea degrade proteins, carbohydrates, fatty acids, amino acids, and hydrocarbons (*5*, *6*, *10*, *62*). Lokiarchaeales, Thorarchaeia, Odinarchaeia, and Heimdallarchaeia are primarily organoheterotrophs with varying capacities to consume and produce hydrogen (*26*). Helarchaeales are proposed to anaerobically oxidize hydrocarbons (*7*, *10*, *63*), whereas Freyarchaeia and Sifarchaeia are predicted to be heterorganotrophic acetogens capable of utilizing methylated amines (*11*, *12*). Hermodarchaeia are proposed to degrade alkanes and aromatic compounds via the alkyl/benzyl-succinate synthase and benzoyl-CoA pathway (*10*). Gerdarchaeales may be facultative anaerobes and utilize both organic and inorganic carbon (*8*). Atabeyarchaeia and Freyarchaeia share several metabolic pathways with new lineages from the Asgard sister-clade TACK (e.g., Brock- and Culexarchaeia) and other deeply branching Asgard lineages. Based on the genomic and metatranscriptomic analyses, we predict that the soil-associated Atabeyarchaeia and Freyarchaeia are chemoheterotrophs that likely degrade amino acids and other carbon compounds. Both encode the EMP Pathway for cellular respiration and the WLP for CO_2_ fixation.

Although Atabeyarchaeia and Freyarchaeia share key central metabolic pathways, they differ in that Freyarchaeia can metabolize compounds such as formaldehyde (C1), glycerol (C3), ribulose (C5), and glucose (C6), whereas, Atabeyarchaeia can only metabolize C1, C3 and C5 compounds (**Fig. 2**). The ability to metabolize C3 and C5 compounds is rare in Asgard archaea. While the entry points into the EMP pathway differ between the two, both exhibit the genetic repertoire necessary for converting carbohydrates into acetate. Both Atabeyarchaeia and Freyarchaeia may also be capable of growth as anaerobic acetogens via acetate production through the WLP. Similar to other Asgard archaea, they have methyltransferase complexes involved in the catabolism of quaternary amines (or yet unknown methylated substrates). Through the use of methylated compounds, they may compete with methanogens and other anaerobic methylotrophic groups that rely on these substrates for methane production. These results align with recent studies suggesting a broader presence of methylotrophic metabolisms among archaea (*10*, *46*, *47*). It also opens up avenues for exploring the environmental impact of these metabolisms, particularly in relation to carbon cycling and greenhouse gas emissions (*64*).

Of particular interest is the predicted metabolic capability of Atabeyarchaeia and Freyarchaeia to degrade aldehydes. Aldehydes in soils come from several sources, including the microbial breakdown of methanol potentially produced from methane oxidation, degradation of plant and animal compounds, and products of industrial combustion and wildfires (e.g., volatile organic compounds). In fact, the California wetland soil that hosts these archaea contain charcoal, likely produced by wildfires. They are also predicted to be capable of growing on glycerol under anaerobic conditions capacity previously undescribed in Asgard archaea. Glycerol may be present in soil by the lysis of bacteria, yeast, and methanogenic archaeal cells that use glycerol as a solute, or by microbial fermentation of plant and animal triglycerides and phospholipids(*65*). The presence of glycerol kinase and the respiratory glycerol-3-phosphate dehydrogenase (GlpABC) in Atabeyarchaeia and Freyarchaeia indicates these archaea might use with glycerol or glycerol-3-phosphate and fumarate as the terminal electron acceptor associated with proton translocation. This finding suggests a broader role for glycerol in Asgard archaeal energy metabolism and points to a possible conservation of this mechanism across different anaerobic environments. Understanding how these archaea metabolize glycerol will enhance our knowledge of their ecological roles and contributions to the carbon cycle in wetland ecosystems. Atabeyarchaeia and Freyarchaeia also produce and consume small organic molecules and H2 that serve as substrates for methane production by methanogens that coexist in wetland soil.

The soil Asgard archaea encodes group 3c [NiFe]-hydrogenase genes, which were shown to be highly expressed *in situ*. Under specific conditions, autotrophic growth is likely supported by H_2_ oxidation via the WLP. The presence of syntenic blocks encoding heterodisulfide reductase complexes, [NiFe] hydrogenases, and ATP synthase suggests a sophisticated apparatus for energy transduction, resembling mechanisms previously characterized in other archaeal groups (*34*). Additionally, our results suggest the existence of an electron bifurcation mechanism in both Asgard archaea lineages, where electrons can be transferred from H_2_ to ferredoxin and an unidentified heterodisulfide intermediate (*26*). Atabeyarchaeia and Freyarchaeia also have membrane-bound group 4 [NiFe]-hydrogenases that likely facilitate the oxidation of reduced ferredoxin generated through fermentative metabolism. However, this complex is novel in that it includes a HL helix on the L-like subunit and two antiporters, neither of which are part of biochemically characterized group 4 respiratory hydrogenases. The functional modeling of these complexes reveals structural congruences with known respiratory enzymes, hinting at a potential for chemiosmotic energy conservation that may be a widespread feature among the Asgard clade. The findings indicate a potential evolutionary connection between hydrogenases and complex I, aligning with the hypothesis that complex I may have evolved from ancestral hydrogenases (*30*, *66*).

These complete genomes provide insight into the unique metabolic pathways of Asgard archaea in soil environments, previously missed in primarily sediment-based descriptions. Of particular interest is the identification of genes encoding enzymes for oxidative stress response in both Atabeyarchaeia and Freyarchaeia, despite their anaerobic nature. The use of selenocysteine in key enzymes may provide another mechanism for dealing with increased oxidative stress. These Asgardarchaeota genomes suggest an adaptation to transient oxidative conditions in soil environments and additional competition for methanogenesis and anaerobic methyltrophy substrates.

## Conclusions

We manually curated three complete genomes for Asgard archaea from wetland soils, uncovering bidirectional replication and an unexpected abundance of introns in tRNA genes. These features suggest another facet of the evolutionary relationship between archaea and eukaryotes. Metabolic reconstruction and metatranscriptomic measurements of *in situ* activity revealed a non-methanogenic, acetogenic lifestyle and a diverse array of proteins likely involved in energy conservation. The findings point to metabolic flexibility and adaptation to the dynamic soil conditions of wetlands. Finally, they contribute to cycling of carbon compounds that are relevant for methane production by coexisting methanogenic archaea.

## Acknowledgments

We thank Basem Al-Shayeb for his contribution to field work and generation of sequence datasets, and Shufei Lei and Jordan Hoff for bioinformatics support. We thank Dr. Chris Greening and Dr. Pok Leung for discussions on archaeal hydrogenases classification. We also thank Adam Panagiotis for the helpful discussion about methyltransferases from methanogens. Lastly, we are grateful to Dr. Luke Oltrogge and Dr. Daniel Gittins for their discussions about the multimeric structures modeled using AlphaFold.

## Author contributions

Brackets denote equal contribution in the author list order. The study was designed by LEVA and JFB. Samples collection and nucleic acid extractions were performed by L.E.V.A., M.C.S, J.F.B., A.C.C., J.W.R. and L.D.S., Metagenomic data was generated by L.E.V.A., M.C.S., R.S., A.C.C., J.W.R., and J.F.B. Genome binning was done by J.F.B., L.E.V.A., M.C.S., A.C.C, and J.W.R. Complete Asgard genome curation was conducted by J.F.B. and L.E.V.A. Phylogenetic analyses were conducted by (KEA and LEVA). Metabolic annotation and analysis done by (LEVA, KEA, and V.D.A.), M.C.S provided knowledge on metabolism of archaea. V.K. performed tRNAs and selenoproteins analysis. D.S. and B.J.B. provided feedback on the study design and methodology. J.F.B., D.S, and B.J.B. provided resources and funding. L.E.V.A. and J.F.B. wrote the manuscript, with significant contributions by K.E.A., V.D.A, M.C.S and input from all authors. All authors read and approved the manuscript.

## Data and materials availability

Prior to publication, the genomes reported in this study can be accessed via https://ggkbase.berkeley.edu/SRVP_asgard/organisms.

