## Supplementary Methods, Tables and Figures for "Asgard archaea modulate potential methanogenesis substrates in wetland soil"

**The PDF file includes:**

**Materials and Methods**

Sample acquisition, nucleic acid extraction and sequencing

We collected soil cores from a seasonal vernal pool in Lake County, California, in October 2018, October 2019, November 2020 and October 2021. Samples were frozen in the field using dry ice, and kept at -80 C until extraction. The Qiagen PowerSoil Max DNA extraction kit was used to extract DNA from 5-10 g of soil, and the Qiagen AllPrep DNA/RNA extraction kit was used to extract RNA from 2 g of soil. Samples were sequenced by the QB3 sequencing facility at the University of California, Berkeley on a NovaSeq 6000. Read lengths for the 2018 DNA samples and the RNA samples were 2x150 bp and 2x250 bp for the 2019-2021 DNA samples. A sequencing depth of 10 Gb was targeted for each of 2018, 2020, and 2021 samples, and 20 Gbp for each of the 2019 samples. A subset of deep soil samples from 2021 were sequenced using PacBio and Oxford Nanopore technologies. For Oxford Nanopore sequencing, the Ligation Sequencing Kit (LSK114) was used to prepare native DNA libraries and the Qiagen Repli-G mini kit and LSK114 were used to prepare amplified DNA libraries. Samples were sequenced on FLO-PRO114M flowcells on the PromethION for 72 hours.

Illumina metagenomic assembly, binning and annotation

Metagenomic sequencing reads were trimmed for adapter sequences and quality using sickle(*69*). The filtered metagenomic reads from 2018 were assembled using the IDBA_UD assembler (*70*) . The reads from 2019-2020 using metaSPAdes(*71*) Contigs greater than 2.5 Kb were retained and sequencing reads from all samples were cross-mapped against each resulting assembly using Bowtie2 (*72*). The resulting differential coverage profiles were filtered at a 95% read identity cutoff and then used for genome binning with MetaBAT2(*73*), VAMB(*74*) and MaxBin2(*75*). The resulting genome bins were assessed for completeness and contamination using CheckM2 (*76*) and were manually curated using taxonomic profiling with GGKBase (www.ggkbase.berkeley.edu). The first iteration of taxonomy was assigned to genome bins with GTDB-Tk v2.3.0 (*77*) and further validated with phylogenetic trees of single-copy marker genes.

We also downloaded all the publicly available Asgard genomes from BV-BRC (www.bv-brc.org) and used CheckM2 to estimate genome completeness.

PacBio metagenomic sequencing and  assembly

Samples from September 9, 2021 from 140cm and 75cm were sequenced using a Sequel II to generate PacBio HiFi reads. Reads were quality trimmed using BBDuk (bbduk.sh minavgquality=20 qtrim=rl trimq=20)(*78*)and assembled with hifiasm-meta(*79*).

Oxford Nanopore metagenomic assembly

Reads were basecalled with Guppy using the dna_r10.4.1_e8.2_sup@v3.5.1 model. Reads were filtered for an average quality >10 and a minimum length of 1kb using fastp (v0.23.2). Adapters were trimmed using porechop (v0.2.4). Branching artifacts caused by multiple displacement amplification were removed by aligning amplified reads to themselves with mappy (v2.24); Reads with non-diagonal self-alignment were removed. Both amplified and native reads were jointly assembled with metaFlye (v2.9)(*80*), long-read polished with medaka consensus (v1.7.1), and short-read polished with Hapo-G (v1.3.1).

Manual genome curation of short-reads genomes and validation using long-reads

The manually curated genomes were de novo reconstructed from high-quality Illumina metagenomic data as described previously(*21*). From soil samples taken at various depths, we recovered draft illumina based MAGs corresponding to Atabeyarchaeia-1, Atabeyarchaeia-2 and Freyarchaeia. The curation process involved the identification and removal of obvious chimeric regions, which were indicated by abrupt changes in GC content or by insufficient Illumina read mapping support. We also corrected sequences in regions with imperfect read alignment, allowing no single nucleotide polymorphisms (SNPs), by mapping reads at a reduced stringency threshold (allowing for up to 3% SNPs). This was followed by manual curation of the consensus sequence, including insertion, deletion, or substitution of individual base pairs. The extension of contig ends was conducted using unplaced Illumina reads. High read coverage was interpreted as indicative of the genomic termini. A genome was deemed complete when it displayed uninterrupted support from Illumina reads. The final assessment of genome completeness was performed by examining the cumulative GC skew and ensuring alignment with known complete genomes from related taxa. The Average Amino Acids of the new genomes was performed using AAI: Average Amino acid Identity calculator tool (http://enve-omics.ce.gatech.edu/aai/) and using compareM (v.0.0.23) with the ‘aai_wf’ at default settings (https://github.com/dparks1134/CompareM) . Replichores of complete genomes were predicted according to the GC skew and cumulative GC skew calculated by the iRep package (gc_skew.py).

Metabolic pathway reconstruction of complete genomes and phylogenetic analysis of key functional genes

We utilized Prodigal (v2.6.3) to predict the genes for Atabeyarchaeia and Freyarchaeia genome by this study(*81*). A suite of databases, including KofamKOALA (v1.3.0)(*82*) and InterProScan (5.50-84.0)(*83*) were combined to begin annotation. Complete genomes were also annotated with

HydDB(*84*), METABOLIC (v4.0)(*85*), PROKKA (v1.14.6), DRAM(*86*) and MEBS (v2.0). Our methods also incorporated a microbial genome annotation workflow as previously described in Undinarchaeota study (*87*).

For each gene/protein of interest, references were compiled by 1) BLASTing the corresponding gene against the NCBI nr database and their top 50 hits clustered by CD-HIT using a 90% similarity threshold, 2) obtaining sequences from the high quality manually annotated UniProtKB/Swiss-Prot/InterPro/PFAM (reviewed) (*catalases*), or 3) using previously published alignment and creating hmm models to extract Atabeyarchaeia and Freyarchaeia sequences (*AOR, catalases,* and *MttB-like homologs*). The final set genes/proteins were aligned using MAFFT v.7.407 (*AOR* and *catalases*)/v7.490 (*ADH* and *MttB-like homologs*), trimmed with trimAl v1.4.rev15 (*AOR, ADH, catalases,* and *MttB-like homologs*), and a phylogenetic tree was inferred using IQTREE v.1.6.6 (*AOR* and *catalases*)/v2.0.7 (*ADH* and *MttB-like homologs*) using automatic model selection. More detailed descriptions for each phylogeny, including model selection are discussed in the supplemental figure captions.

### Identification of [NiFe]-hydrogenase and phylogenetic analyses

We extracted [NiFe]-hydrogenase sequences from complete genomes as well as publicly available Asgardarchaeota genomes from NCBI and BCWT using custom Hidden Markov Models (HMMs). To confirm the accuracy of the candidate [NiFe]-hydrogenase sequences, we checked the identified sequences by looking at nearby genes and ensuring the presence of essential hydrogenase accessory genes and confirmed the preliminary classification using HydDB(*84*). We combined our [NiFe]-hydrogenase sequences with those from the HydDB database. Then, we aligned all the sequences using MAFFT (parameters:--localpair --maxiterate 1000). After alignment, we looked for the N-terminal and C-terminal CxxC conserved motifs. We then used TrimAl v (parameters: -gt 0.5) to clean up the alignment, removing parts where more than 50% of the sequences had gaps. We used IQ-TREE version 1.6.12 with the best-fit model according to Bayesian information criterion (BIC) and ultrafast 1000 bootstrap method to estimate support values for the tree branches. The output tree was visualized using ItoL and the subgroups were classified based on known HYDB references(*84*).

### Identification of Selenocysteine machinery and Selenoproteins, tRNA and intron predictions

The Sec machinery (pstK, EFsec, SecS, and SPS) was identified using HMMER v3.3.1 (*88*) via hmmscan against the TIGRFAM v15.0 database (*89*) and also confirmed via Selenoprofiles v4.4.8 (*90*), using the -p machinery option. Selenoprofiles v4.4.8 and the Seblastian- SECIS3 software (*91*) was accessed to identify potential selenoproteins, which were manually curated via alignment to known selenoproteins as previously reported (*92*). SECIsearch3 was used to identify SECIS for selenoproteins.

For identification of predicted pre-tRNA, we used tRNAscan-SE v.2.0.12 (default settings and -A for archaea model) (*93*). The R2DT software (*94*) was used to predict and visualize tRNA secondary structure, which also led to identification of additional introns that were not detected by tRNAscan. We identified Sec tRNA with the Secmarker v0.4 webserver (*95*) with a minimum Infernal score of 35.

Metatranscriptome sequencing and analysis

A subset of the deep SRVP soil samples (60cm, 70cm, 80cm and 100cm) were used for metatranscriptomics. Asgard genomes were first dereplicated using derep v 3.4 (*96*), resulting in a set of representative genomes. A Bowtie2 index was constructed from the dereplicated genomes to expedite the sequence alignment process. Post alignment, a custom-built Python script(*97*) was utilized to scrutinize the paired reads from the alignment (BAM) files and deliver statistical insights on gene expression. This script initiates by creating a Pysam 'AlignmentFile' object, enabling a systematic traversal through the '.bam' files by parsing the complex binary alignments. It then proceeds to iterate over each gene listed in a predefined set. For each gene, the script tallies the reads that satisfy user-defined thresholds for Average Nucleotide Identity (ANI) and Mapping Quality (MAPQ). These thresholds, designed to uphold a high level of alignment quality and sequence identity, are critical in enhancing the reliability of the data. Upon the completion of this process, the script outputs the filename, the gene name, and the count of reads that conform to the defined parameters. This pipeline permits an in-depth investigation of RNA hits within the dereplicated genomes, thereby illuminating gene expression profiles across diverse environmental scenarios.

To quantify the number of reads per gene, we further utilized a Python script (filter_counts.py). This script operates based on gene predictions produced by the Prodigal software for the genomes of interest. With the required gene predictions secured through a rerun of Prodigal, the script was set in motion on the BAM files. The command executed was as follows: python filter_counts.py -q 10 -m 0.97 derep_genomes.faa. This command sets the minimum MAPQ score (-q 10) and the minimum ANI (-m 0.97) to ensure high-quality alignments and a high degree of sequence identity. This adherence to stringent criteria bolsters the reliability and robustness of our analytical process.

Mapped mRNA reads were standardized by the calculation reads per kilobase of transcript per million reads mapped (RPKM), which allows for the direct comparison of the level of transcription of all genes in the de novo assembly (**table S4**). RPKM is calculated in Rockhopper as: RPKM=((number of mapped reads/length of transcript (gene) in kilobase)/million mapped reads))(*98*).

Abundance and distribution of archaea

The dereplicated Atabeyarchaeia, Freyarchaeia and other archaeal genomes present in the 36 metagenome samples were mapped to all the metagenome reads independently using BBMap(*78*) (parameters: nodisk=t pigz=t unpigz=t ambiguous=random). We set the minimum identity for mapping to 0.9 and handled ambiguous mappings randomly. The percentage of relative abundance of each genome, and the unmapped read percentage values were calculated using coverM (parameters: -m relative_abundance --min-read-percent-identity 95).

Phylogenetic analyses of Asgard genomes

We used three different sets of markers for the phylogenetic analysis. 47 arCOGs (Archaeal cluster of orthologous genes), a subset of previously described marker set (*99*), were extracted with hmmsearch (v3.1b2) from the 9 MAGs in this study, 303 additional Asgard MAGs, and 36 TACK publicly available MAGs. Korarchaeales MAGs were excluded from the analysis due to the hyperthermal-sensitive marker genes (*4*). We chose to exclude arCOG01183 from the previous set as it was found in less than 50% of the MAGs. mafft (v7.310) and BMGE (v1.12) were used to align and trim the concatenated alignments. Maximum likelihood phylogenetic trees were generated using IQ-TREE (v1.6.1) to test various models and topologies to obtain bootstrap values and LG+F+R10 model was selected (**Fig. 1C**). Additionally, we compared this phylogeny with a set of 37 single-copy marker genes from Phylosift (v1.0.1)(*100*) (Darling et al., 2014). The sequences were manually accessed, aligned with mafft auto (v7.490), trimmed with BMGE (v2.0), and a maximum likelihood phylogeny generated with IQ-TREE (v2.0.7), using the LG+F+R10 model (**fig. S24**). All marker sets resulted in the same phylogeny for the 9 added MAGs. 16S ribosomal proteins were extracted from 439 Asgard and 39 TACK MAGs with barrnap 0.9, using options --kingdom arc --lencutoff 0.2 --reject 0.3 --evalue 1e-05. Sequences manually curated, aligned with mafft auto (v7.490) and masked with Geneious Prime, and ran with IQ-Tree (v2.0.7). 16S phylogeny separates Atabeyarchaeia MAGs from other Asgard clades (**fig. S2**). Amino acid identity (AAI) was determined with CompareM (v0.0.23) to distinguish taxonomic level. We assigned numbers from 1-9 as candidate groups within this new Asgardarchaeota class.

Identification and analysis of genomic ESPs

PSI-BLAST was used to query the Asgard proteomes against the AsCOGs (Asgard database) and arCOGs databases (*3*). Hits were only considered if the absolute difference between the subject and query sequence was less than 75% of the length of the query sequence. Hits were then dereplicated by taking the best hit for each query.

Protein structure prediction using AlphaFold-multimer and ColabFold

Structural predictions for group 4 [NiFe]-hydrogenases, CoxLMS, MtrA, and MtrAH fusion proteins were performed using ColabFold, incorporating AlphaFold-multimer capabilities (*101*). These predicted structures were then visualized and aligned using ChimeraX (*102*), with corroborating reference structures obtained from the Protein Data Bank (*103*). This structural modeling was integral in reinforcing our phylogenetic and metabolic inference.

Nomenclature, etymology, and proposal of type material

Based on the findings of our research, we propose a new taxonomic nomenclature for the identified Asgard archaea organism. The chosen nomenclature encapsulates various facets that resonate with the organism's nature and its ecological habitat. The term 'Atabey' is a homage to the Mother Goddess, often referred to as the Earth Mother in the Taíno mythology, symbolizing fertility, abundance, and the nurturing aspects of nature. e suggest the taxonomic hierarchy of class 'Ca. Atabeyarchaeia', order 'Ca. Atabeyarchaeales', family 'Ca. Atabeyarchaeaceae', and genus 'Ca. Atabeyarchaeum'. We submit the complete genomes “Atabeyarchaeia-1” and “Atabeya-2” as well 6 near complete genomes as type material for this new group.

### Supplementary Text

#### Extended Metabolism:

**Modified EMP Pathway:**
Atabeyarchaeia and Freyarchaeia use the Embden-Meyerhof-Parnas (EMP) glycolytic pathway, producing ATP through fermentation of acetyl-CoA to acetate. The entry point into the pathway is F6P for the Atabeyarchaeia genomes and glucose for Freyarchaeia. In addition to the different entry points, they utilize different enzymes to produce pyruvate. Atabeya encodes the oxygen-sensitive reversible enzyme, pyruvate phosphate dikinase (ppdK; IPR010121); whereas Freya encodes unidirectional pyruvate water dikinase/phosphoenolpyruvate synthase (pps; IPR006319) and pyruvate kinase (pk; IPR001697), producing phosphoenolpyruvate and pyruvate, respectively. The pyruvate phosphate dikinase found in Atabeya is proposed in archaea to be an ancestral version of components of the glycolytic pathway in anaerobic eukaryotes (*104*), **fig. S13**). Freyarchaeaia encoding a complete EMP Pathway supports previous descriptions of Freyarchaeaia MAGs from sediments and other Asgard archaea, including Lokiarchaeia, Hermodarchaeia, Thorarchaeia, Odinarchaeia, Sifarchaeia, and Heimdallarchaeia(*10*, *12*). Less common among the described Asgard is starting at F6P, lacking the traditional enzymes for the degradation of glucose at the start of the EMP Pathway. In this capacity, Atabeyarchaeia more closely resembles the pathway's description for Helarchaeales, an order of Lokiarchaeia(*7*). Helarchaeales have not been shown to rely on simple sugars, but instead, utilizes alkanes as an alternative carbon source, connecting glycolysis to a partial TCA cycle. From the two complete Atabeyarchaeia genomes, we were able to map transcripts from G6P to 2PG and PEP to acetate. Atabeyarchaeia transcripts supported metagenomic data, connecting the TCA cycle (oxaloacetate) to EMP Pathway (PEP) as both genomes transcribed phosphoenolpyruvate carboxykinase. Freya transcribed 3PG to 2PG and PEP to acetate. Like Atabeyarchaeia, Freyarchaeia transcribed enzymes connecting malate and oxaloacetate in the TCA cycle to EMP and a portion of RuMP/NO-PPP, connecting H6P to F6P.

**Partial TCA Cycle:** Atabeyarchaeia and Freyarchaeia resemble previous descriptions of Asgardarchaeota, encoding only partial TCA cycle, connecting this pathway to the EMP Pathway and amino acid degradation (i.e., glutamate). The main difference between lineages in this pathway is that Atabeyarchaeia encodes fumarate hydratase, catalyzing the reversible reaction of fumarate to malate. The only portion of the TCA cycle transcribed in Atabeyarchaeia and Freyarchaiea is 2-oxoglutarate/2-oxoacid ferredoxin oxidoreductase, producing reducing power in NADH.

**Hydroxyglutarate Pathway:** Freyarchaeaia and Atabeyarchaeia appear to ferment amino acids via the partial hydroxyglutarate pathway. Of the three main hydroxyglutarate pathway steps, both lineages encode the first reversible NADH-dependent reduction of 2-oxoglutarate to 2-hydroxyglutarate (step 22 in Figure 2), connecting amino acid fermentation to the partial TCA cycle and therefore production of ATP and NADPH or NADH reducing equivalents. Only Freyarchaeia has the enzymes for the dehydration of hydroxyglutaryl-CoA to glutaconyl-CoA (HgdAB) and both groups lack the enzyme necessary to decarboxylate glutaconyl-CoA to crotonyl-CoA (*105*). The encoded hydroxyglutarate pathway enzymes (HgdAB, GdhA, PHGD, GctAB) may produce H+, ammonium, acetate and hydroxyglutaryl-CoA. Assuming there is a mechanism to convert glutaconyl-CoA to crotonyl-CoA, fermentation of crotonate would produce NADH, $FADH_2$ and acetyl CoA. This pathway links amino acid fermentation to central metabolism and production of ATP and is consistent with descriptions of anaerobic amino-acid degradation in the last Asgard common ancestor (*4*, *9*).

**Reductive Glycine Pathway (rGlyP):** We identified genes predicted to be involved in the anaerobic oxidation of glycerol and the glycine cleavage system (GCS), which is part of the reductive glycine pathway (rGlyP). All three genomes encode glycerol-3-phosphate dehydrogenase complex (*glpABC*), a putative glycerol kinase (*glpK*), and glycerol dehydrogenase (*gldA*), as well as, P, T and H proteins of GCS. Sequences from Atabeyarchaeia and Freyarchaeia are phylogenetically distinct from characterized glycerol-3-phosphate dehydrogenase subunit A (*glpA*), however Freyarchaeia clusters with Halobacteriales known to metabolize glycerol (*106*) (**table S7**). Reverse GlyP (rGlyP) has been identified in other archaea but this pathway in Atabeyarchaeia and Freyarchaeia is unique in that it resembles *E. coli*, lacking the L-protein (dihydrolipoyl dehydrogenase), reducing NAD+ to NADH (*46*, *107*). Despite the discussion of glycine metabolism in archaea little is known about the the rGlyP in Asgard archaea (*9*, *11*). Yet the ancestral reconstructions suggest the glycine cleavage system was present in the Asgard archaeal ancestor, which supports the suggested deeper phylogenetic branching of these two soil Asgard clades (*4*).

**Hydrogenases:** Atabeyarchaeia and Freyarchaeia generate energy through various NiFe hydrogenases, incomplete electron transport chain, and glycerol respiration. Both clades harbor Group 4g and Group 3c [NiFe]-hydrogenases. Membrane-bound Group 4 [NiFe] hydrogenase (*108*) has previously only been reported in Hermod-, Heimdall- and Odinarchaeia. As Atabeyarchaeia and Freyarchaeia lack methyl coenzyme-M (*mcrABC)* genes, HdrA2B2C2 may function bidirectionally, facilitating both hydrogen oxidation and the formation of a bifurcating complex, coupling the presence of Group 3c [NiFe]-hydrogenase (*MvhADG*) genes. Previous descriptions of Asgard and TACK lineages have suggested Group 3c [NiFe] hydrogenase: heterodisulfide reductase-linked hydrogenases are involved in energy conservation through electron bifurcation (*1*, *47*). Similar to other TACK lineages, both Atabeya genomes have Group 3b [NiFe]-(sulf)hydrogenase, coupling oxidation of NADPH to fermentative production of $H_2$ or sulfhydrogenase activity, reducing elemental sulfur to hydrogen sulfide previously reported in *Pyrococcus furiosus (109)*. The addition of Group 3b [NiFe]-hydrogenase (*HydABCD*) in Atabeyarchaeaia enables additional hydrogen formation, which is absent in Freyarchaeia (*26*, *110*, *111*). The presence of [NiFe]-hydrogenase Group 3b and 3c indicates that these archaea potentially use $H_2$ as an electron donor, or more likely they generate fermentation capacity, as suggested by others(*34*, *112*). A membrane-bound succinate dehydrogenase complex and an A-type ATP synthase complex are encoded in Freyarchaeia and Atabeyarchaeia. Both groups are capable of transferring electrons from glycerol-3-phosphate generated from the reduction of menaquinol to reduce fumarate to succinate as the final electron acceptor in the cell (*113*).

**Alcohol Dehydrogenases and Butanol Oxidation:** Atabeyarchaiea and Freyarchaeaia complete genomes encode iron-containing alcohol dehydrogenase (Fe-ADH)-like enzymes, which likely participates in both fermentation and production of acetyl-CoA. This capacity is also predicted in Hel-, Njord-, and Lokiarchaeales, as supported by phylogenetic analysis **fig. S22**. In particular, the Freyarcheaia complete genomes contains a potential NADPH-dependent butanol dehydrogenase (BDH) part of the reversible butanol oxidation/pyruvate fermentation to butanol pathway, a prevalent fermentation product in both wetland sediments and terrestrial carbon cycling (**fig. S23**). Butanol can be used as a substrate for anaerobic methanogenesis, converting butanol into butyrate, acetate, and methane, sequentially.

**Aerobic Carbon-Monoxide Dehydrogenase (CoxLMS):** We also identify putative genes for aerobic carbon-monoxide dehydrogenase (CoxLMS) and cofactors within Freyarchaeia. The putative *CoxL* forms a monophyletic group with other archaea, suggesting a potential capacity for carboxydotrophy or the utilization of alternative substrates in the presence of oxygen, such as aldehydes or purines, as a member of the aldehyde oxidase superfamily (**fig. S12**)

Besides the bifunctional carbon monoxide dehydrogenase that is part of the WLP and generates acetyl-CoA, Freyarchaeia also encodes putative genes for aerobic carbon-monoxide dehydrogenase (CoxLMS) and cofactors which could oxidize CO as an additional electron donor, suggesting a potential capacity for carboxydotrophy. However, phylogenetic analysis shows that the CoxL from Freyarchaeia belongs to a monophyletic group and clusters with other uncharacterized archaeal CoxL and does not cluster with biochemically characterized form I CoxL suggesting the potential use of other substrates.

**Biosynthesis of Coenzymes and other metabolic precursors:** Other operons suggest that Atabeyarchaeia and Freyarchaeia can self-synthesize MoCF (Molybdenum Cofactor, Molybdopterin), a cofactor enabling the functionality of a wide spectrum of enzymes (i.e., xanthine dehydrogenase superfamily), generating NADH or NADPH. Both lineages also have the enzyme for the biosynthesis of coenzyme A (*pok*), a precursor in various metabolic processes, including the previously discussed TCA cycle and amino acid metabolism. Additionally, Freyarchaeia has enzymes for the biosynthesis of coenzymes B (*leuABCD*), M (*thrC*, *comABCDE*), and F420 (*cofCDEGH*) not encoded in Atabeyarchaeia. The Asgardarchaeota sister-lineage TACK has been shown to have similar enzymes for cofactor biosynthesis (*47*).

**Metatranscriptomics:** Transcriptomic data indicates *in situ* expression acetogenic and energy conservation pathways in both Atabeyarchaeia and Freyarchaeia complete genomes, including EMP glycolysis/gluconeogenesis (**fig. S13**), WLP (**fig. S9**), beta oxidation, formaldehyde oxidation, RuMP (**fig. S15)**, Group 3c [NiFe] Hydrogenase, ATP synthase (**Figure 2**, step 82), NADH-quinone oxidoreductase (**Figure 2**, step 85), and molybdopterin biosynthesis (**table S87)**. We also identified transcription for environmental and stress response, including nickel, arsenite, magnesium, iron, and copper transporter , heat shock proteins, catalase, and superoxide dismutase (**fig. S20**).

Both Atabeya-1 and Atabeya-2 had more than 10 copies of sugar and amino acid transporters, and GT CAZYmes for glycosylation activities, providing the substrates for the fermentation of sugars and amino acids. In Atabeya-1, we identified another highly transcribed (>10 transcript) gene (*pckA*), potentially replenishing the phosphoenolpyruvate (PEP) pool for glycolysis/gluconeogenesis. The eukaryotic signature protein, Lokiactin, was highly expressed in Atabeyarchaeaia, being the most expressed gene in Atabeya-1. Atabeya-2 had more than 10 copies for the Group 3b [NiFe] hydrogenase components not found in Freyarchaeia, *mtrA*, *aor*, both a peptidase inhibitor (I87) and an asgparagine peptide lyase (N11), and amino acid metabolism genes (*trpB, kce, iorA*) ( **table S7**).

In Freyarchaeaia, we identified more than 10 transcript copies of beta-glucosidases, cytoplasmic/cytoplasmic membrane CAZYmes (AA0, CBM50, GH1, GH3, GT3, GT7, and GT66), asparagine peptide lyase (N11), beta oxidation, *aor*, and an intermediate step of the WLP H4MPT branch (*mer*). Environmental stress responses were among the most highly transcribed, especially the catalase-peroxidases not found in Atabeyarchaeia, potentially detoxifying hydrogen peroxide with methanol to produce formaldehyde, superoxide dismutase, small heat shock proteins, and copper detoxification transport **(fig. S20, table S7)**.

**Fig. S1** Maximum-likelihood tree, inferred with IQtree and the best-fit LG+C20+F+G model, using a concatenated set of RP15. Ultrafast bootstrap support values of >90 are shown.

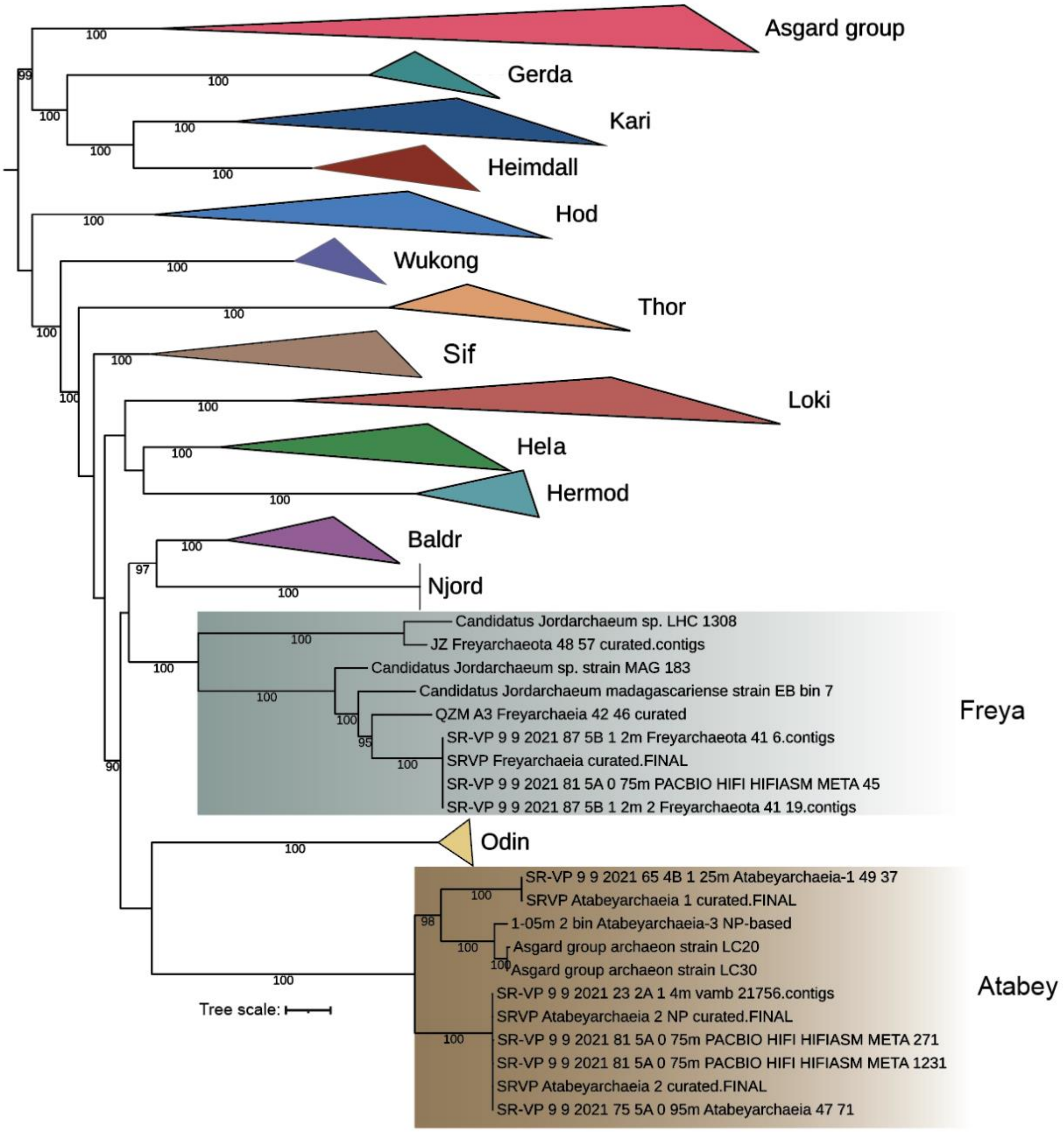

**Fig. S2** Maximum-likelihood tree, inferred with IQtree and the best-fit GTR+F+R6 model, using 16S ribosomal proteins from Asgard and TACK. Ultrafast bootstrap support values of >80 are shown.

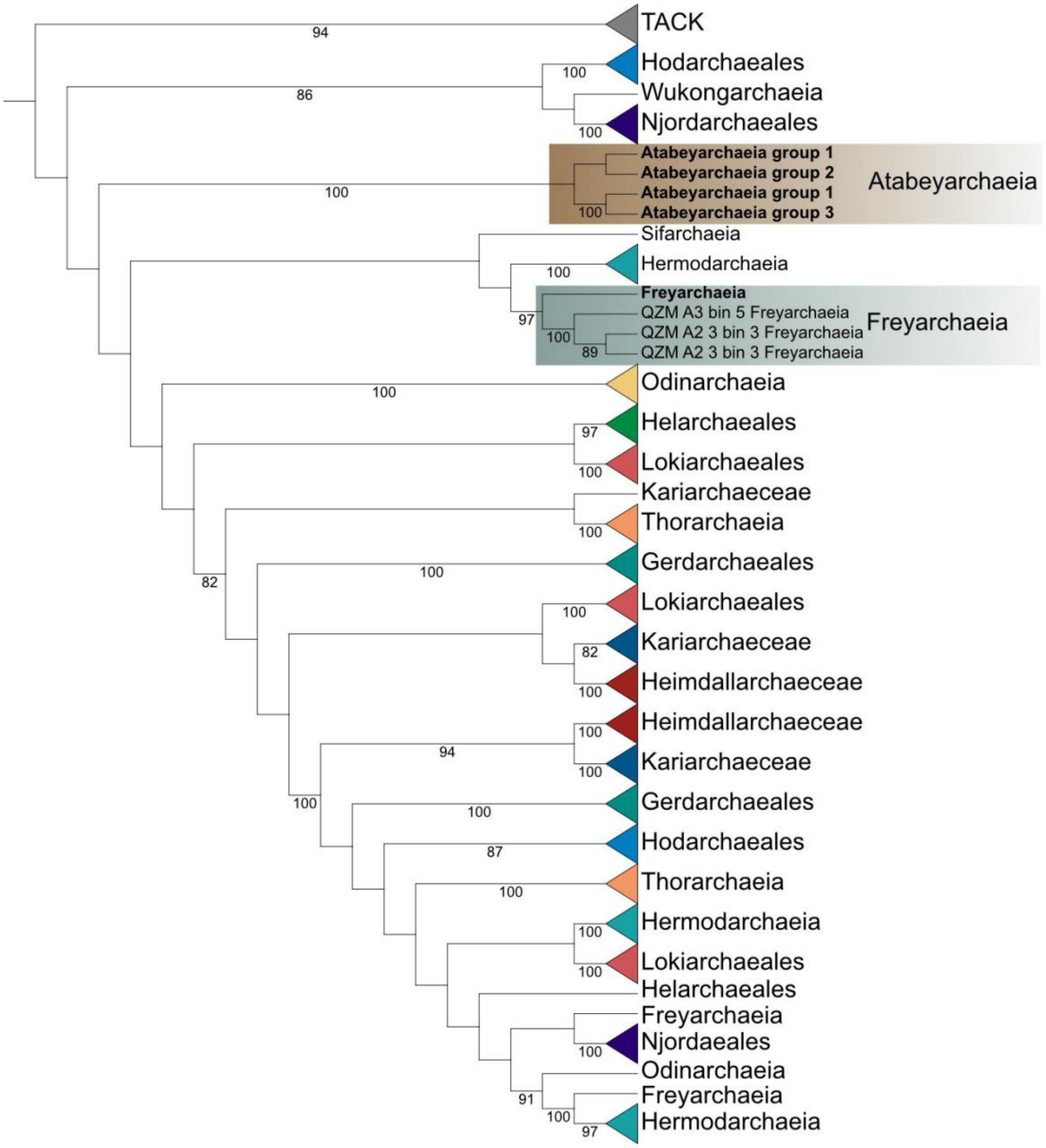


**Fig. S3** Bidirectional replication in Atabeyarchaeia-2 and Freyarchaeia complete genomes. The GC skew is shown as a gray plot and the cumulative GC skew is overlain (green line).

SRVP Atabeyarchaeia-2

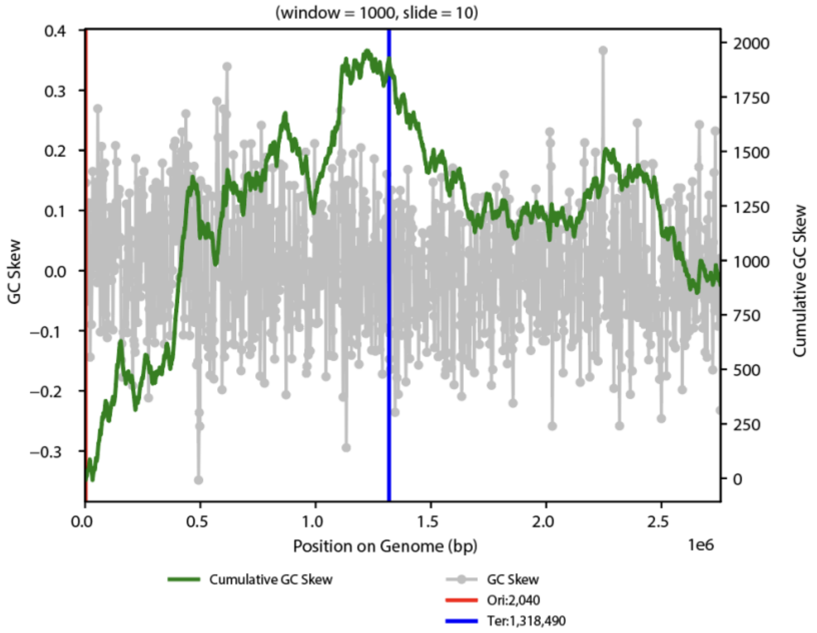


SRVP Freyarchaeia

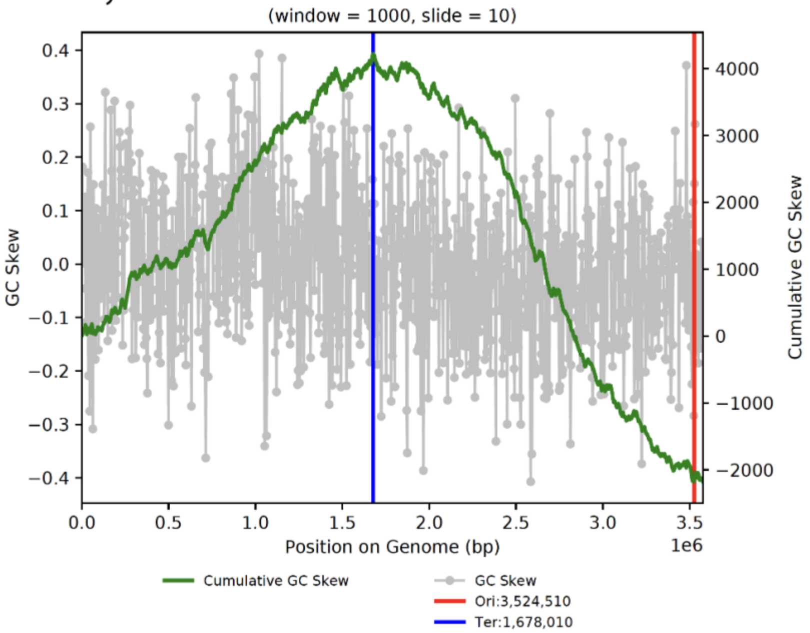

**Fig. S4** Confirmation of complete Atabeyarchaeia genome architecture reported based on Illumina assemblies. Overall topology of genomes for Atabeya-2 PacBio assembled circular genome, Illumina manually curated genome and assembled nanopore genome.

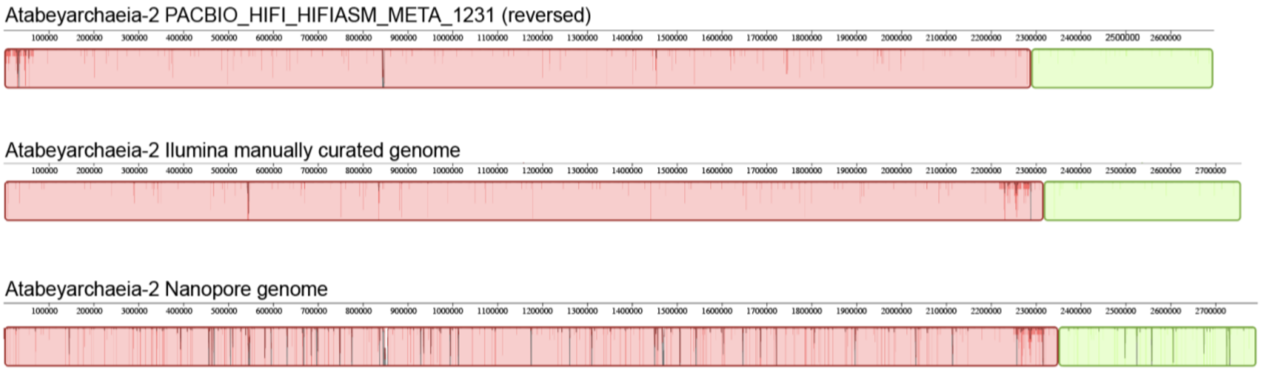

**Fig. S5** Nanopore assembled circular contig revealed another Atabeyarcaheia group-3.

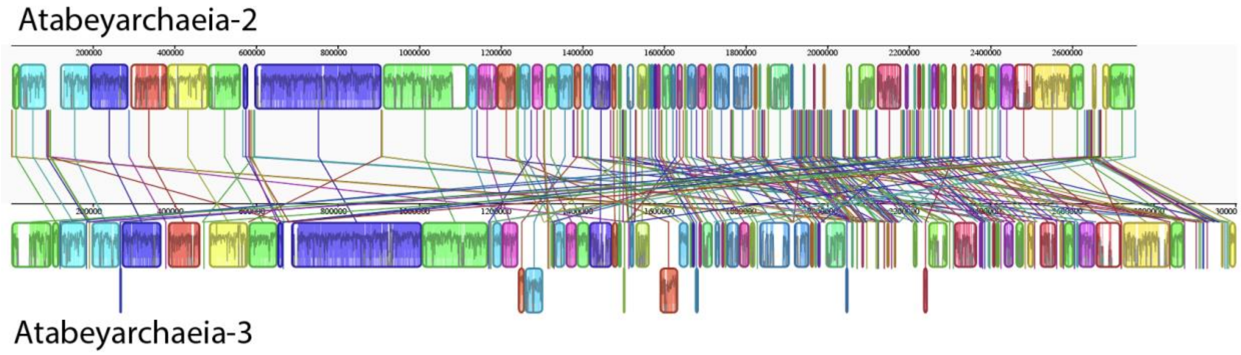

**Fig. S6** Distribution of eukaryotic signature proteins (ESPs) present in Atabeyarchaeia and Freyarchaeia genomes used in this study. ArcCOGs functional categories are **J**, Translation, ribosomal structure, and biogenesis; **L**, Replication, recombination, and repair. **K**, Transcription; **L**, Replication, recombination, and repair. **M**, Cell wall/membrane/envelope biogenesis; **R**, General function prediction only; **S**, Function unknown. **O**, Post-translational modification, protein turnover, and chaperones; **H**, Coenzyme transport and metabolism.

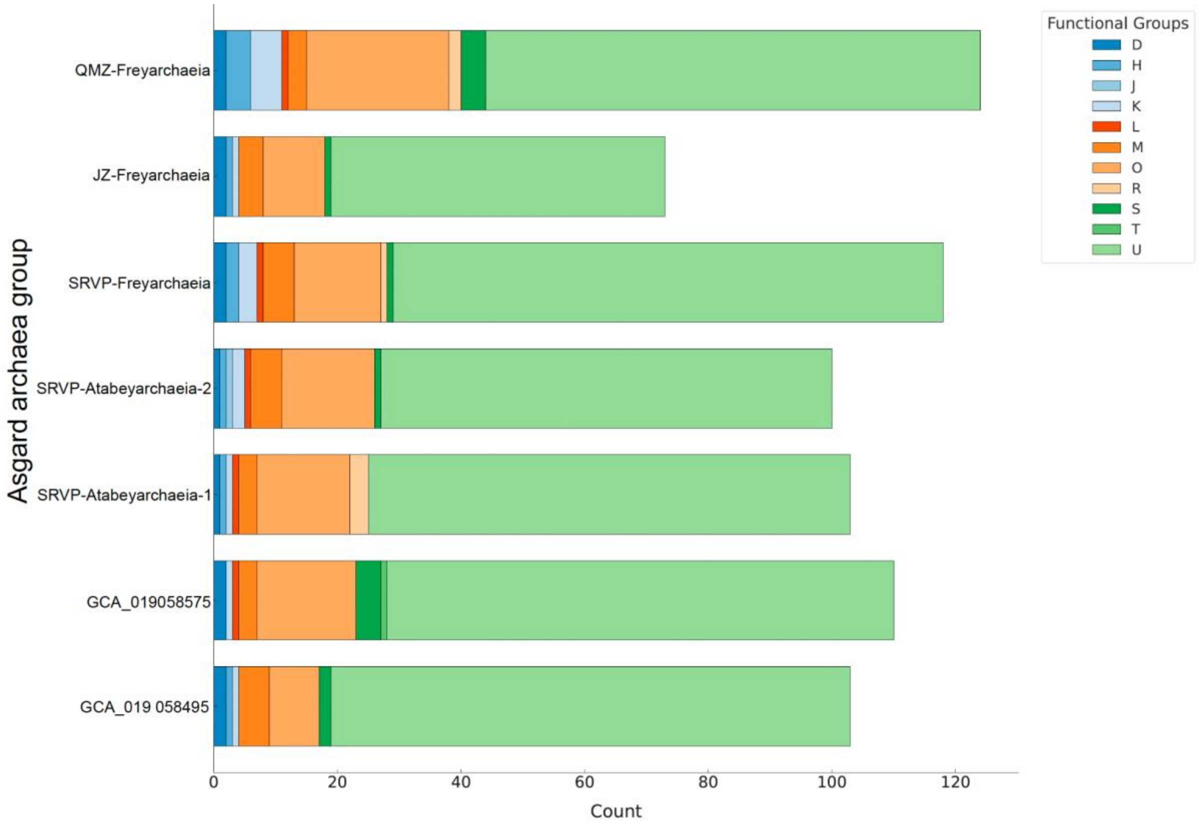

**Fig. S7 Structural superimposition of characterized complexes. A.** AlphaFold models of [NiFe]-hydrogenase module and the proton-translocating membrane module where each candidate subunit is represented by a different color based on the best subunit matched. **B.** Superimposition of the formate hydrogenlyase complex from E. coli. **C.** Superimposition of the Mrp antiporter complex. **D.** Superimposition of the respiratory complex I hydrogenase from *Pyrococcus* , with hydrophobic regions marked in orange to denote areas rich in hydrophobic amino acids. **E.** Model of the complex with surface electrostatic potential coloration: blue for positively charged regions, red for negatively charged areas, and white for neutral zones. This color scheme elucidates the charge distribution on the protein's surface, providing insights into potential interactions with other biomolecules or ions

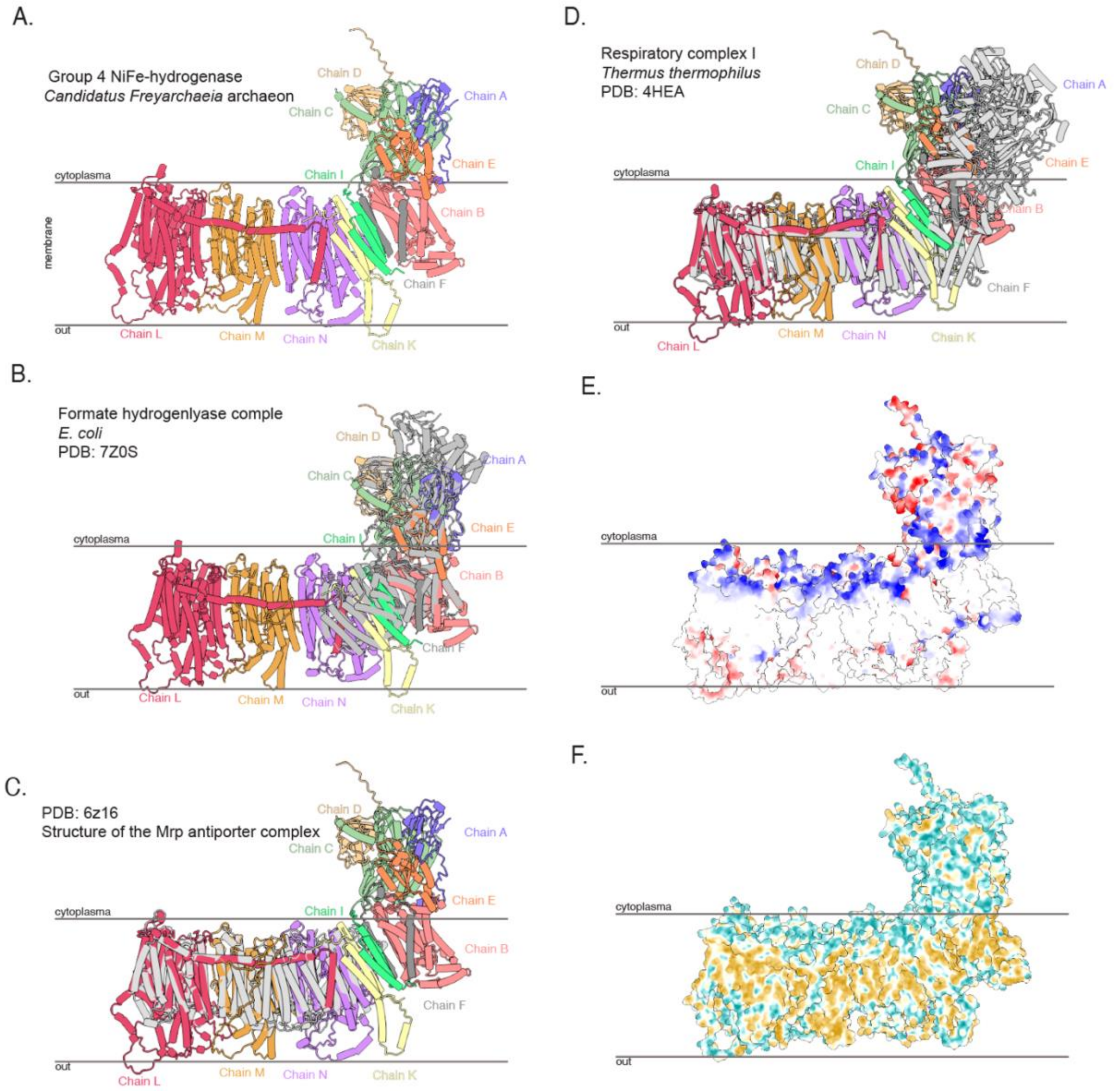

**Fig. S8** Maximum-likelihood phylogenetic reconstructions of the large subunit of group 3 [NiFe]-hydrogenases. (A) Group 3c (b) Group 3b The trees were generated with IQ-TREE v2.0.7, model LG+F+R10 was chosen according to BIC. Ultrafast bootstrap support values of >90 are shown.

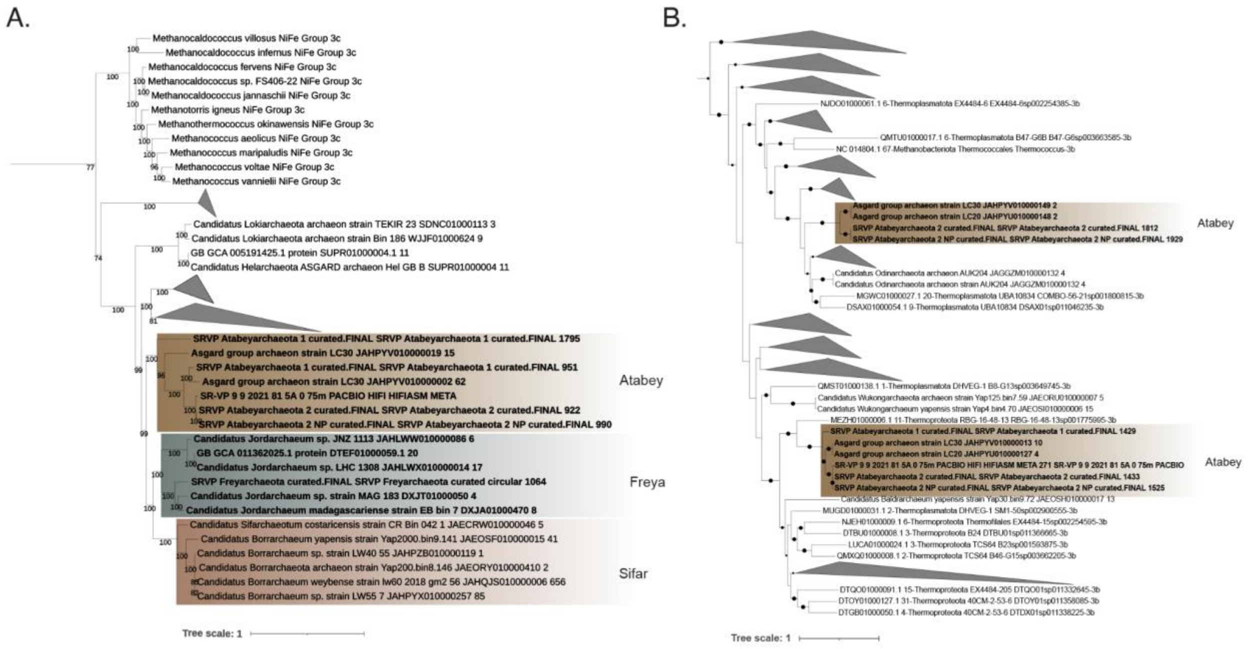

**Fig. S9:** Genes and transcripts for the Wood-Ljungdahl Pathway (WLP). The gene numbers in circles correspond to those in Figure 2 and supplementary table 8. Red arrows indicate that transcripts were detected. Step 115 is unique to Atabeyarchaeia. Created using BioRender.com.

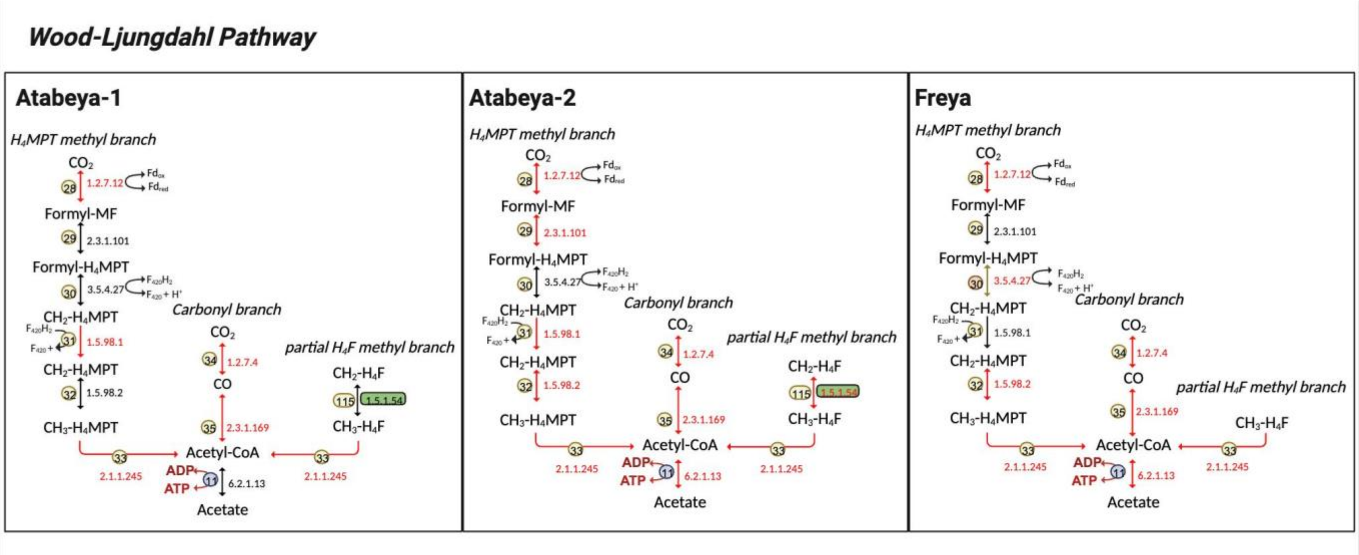

**Fig. S10: Non-Pyl trimethylamine methyltransferase homologs are widespread in Asgardarchaeota** Maximum likelihood phylogeny of non-Pyl trimethylamine methyltransferase A subunit homologs (MttB). We identified two Atabeyarchaeia and two Freya MttB-like in the complete genomes, which were aligned with 800 references(*114*) and 25 additional blastp hits. The sequences were aligned and trimmed with MAFFT v7.505 and trimAl v1.4.rev15. The tree was generated with IQ-TREE v.1.6.12, model LG+C20+R+F. Ultrafast bootstrap support values of >80 are shown.

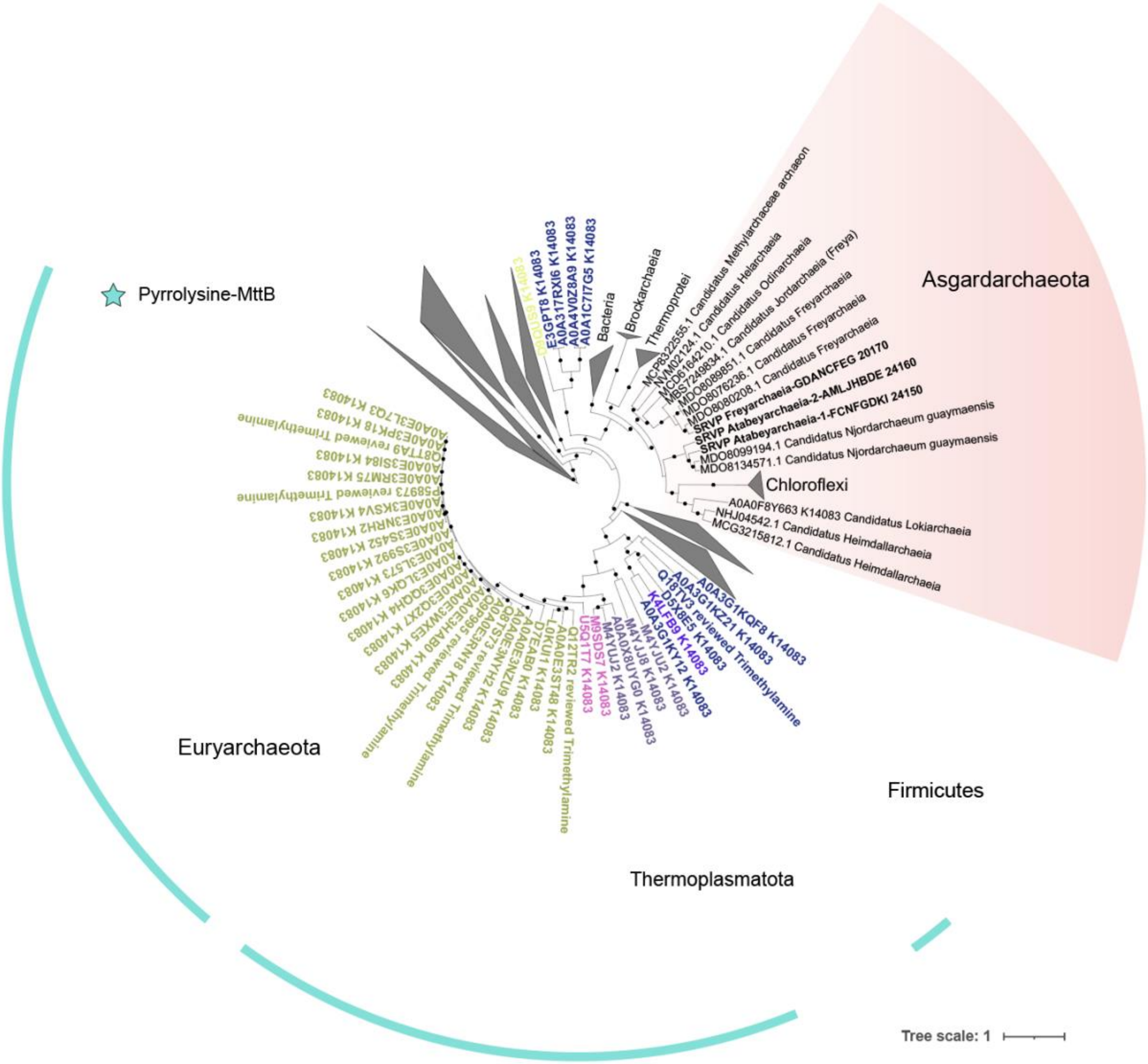

**Fig. S11: Non-Py MtbC dimethylamine-specific corrinoid protein (MtbC) homologs** Maximum likelihood phylogeny of Non-Py dimethylamine-specific corrinoid protein homologs (MtbC and MtbC-like). We identified two Atabeyarchaeia and two Freyarchaeia MtbC-like in the complete genomes, which were aligned with a subset of 10 references based on protein identity from Methanogenic archaea and 25 additional blastp hits. The sequences were aligned and trimmed with MAFFT v7.505 and trimAl v1.4.rev15. The tree was generated with IQ-TREE v.1.6.12, model LG+C20+R+F. Ultrafast bootstrap support values of >80 are shown.

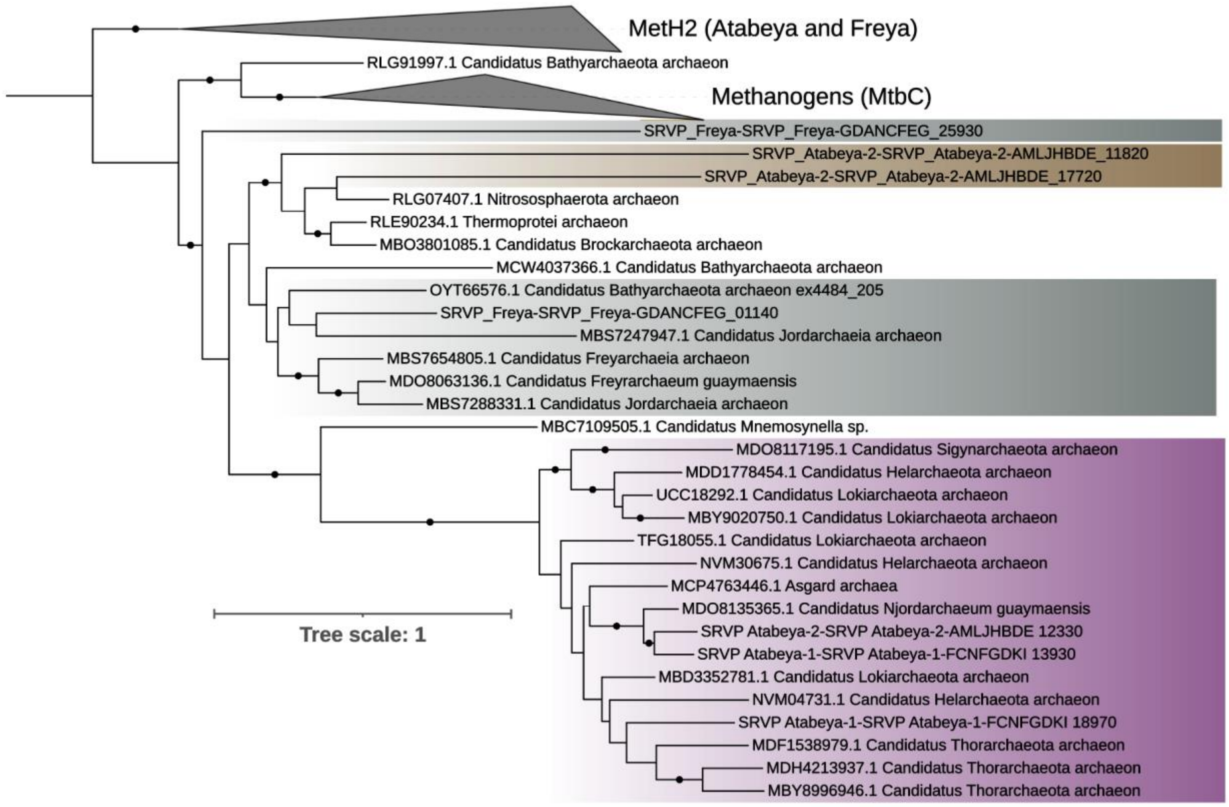

**Fig. S12**: Novel archaeal aerobic carbon monoxide dehydrogenase in Asgard archaea. **A.** Maximum likelihood phylogenetic tree of the aerobic carbon monoxide dehydrogenase large subunit (CoxL) sequences identified in Asgard archaea. It includes a comparison with reference sequences from established categories such as bonafide CoxL type I and II, halobacterial CoxL, as well as bacterial xanthine dehydrogenase (XDH), and aldehyde oxidases from eukaryotes as well as Asgard archaea. This phylogeny elucidates the evolutionary relationship and potential functional diversification of Coxl within these groups. **B**. The panel presents the genetic architecture of cox genes genes with gene clusters color-coded based on Pfam domain classifications and gene annotations. This organization highlights the structural variability and potential regulatory elements within the genomic context of these archaeal enzymes. Ultrafast bootstrap support values of >80 are shown.

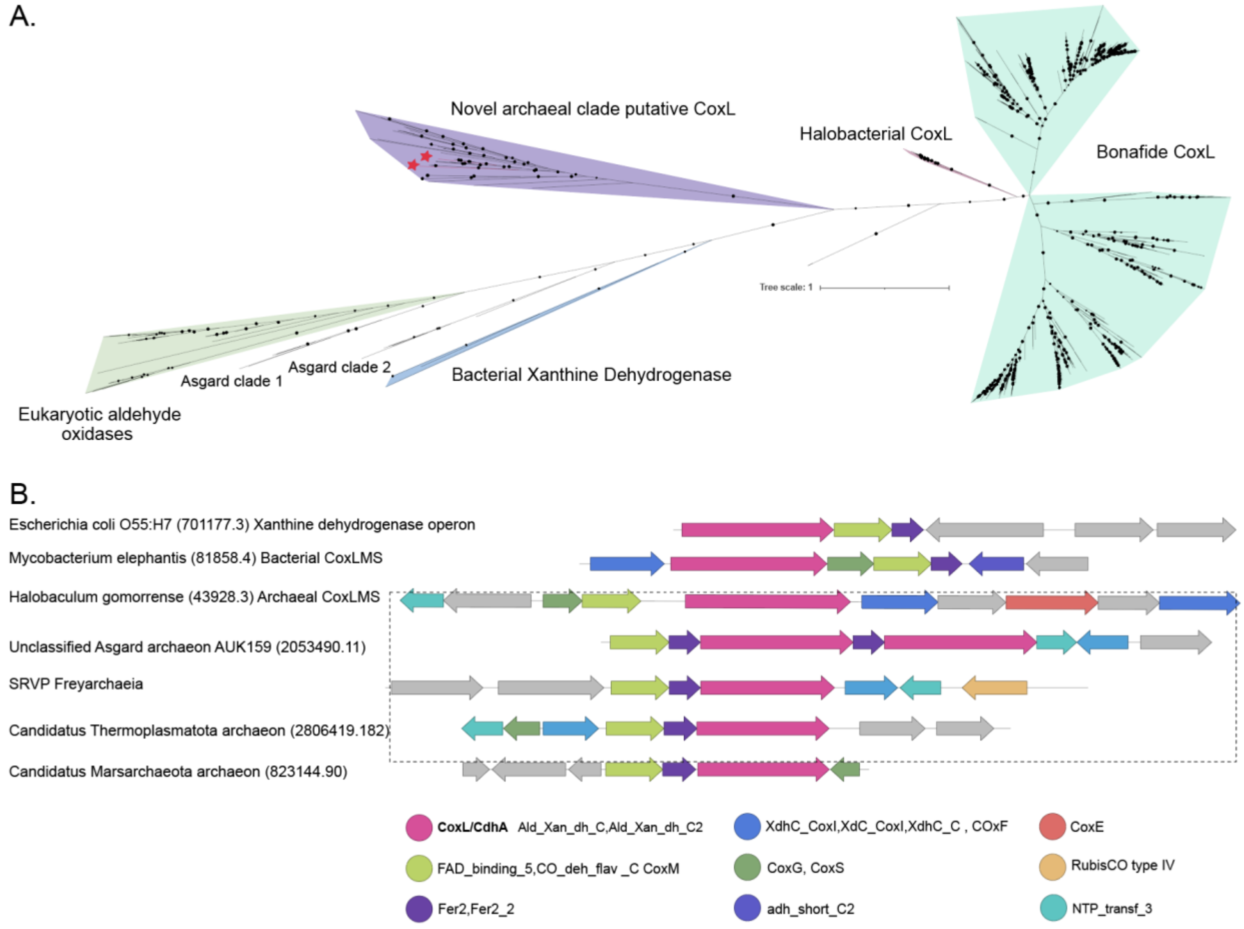

**Fig. S13:** Transcriptome map of the archaeal modified Embden-Meyerhof-Parnas (EMP) Pathway for Atabeyarchaeia (Atabeya-1 and Atabeya-2) and Freyarchaeia (Freya). The numbers correspond to those in Figure 2 and Supplementary Table 7. Red arrows indicate mapped transcripts,the rectangles show the number of transcripts, dashes around EC numbers show oxygen-sensitive enzymes, and the green boxes behind the EC numbers indicate lineage-specific reactions for Atabeya genomes, matching Figure 2. Created using BioRender.com.

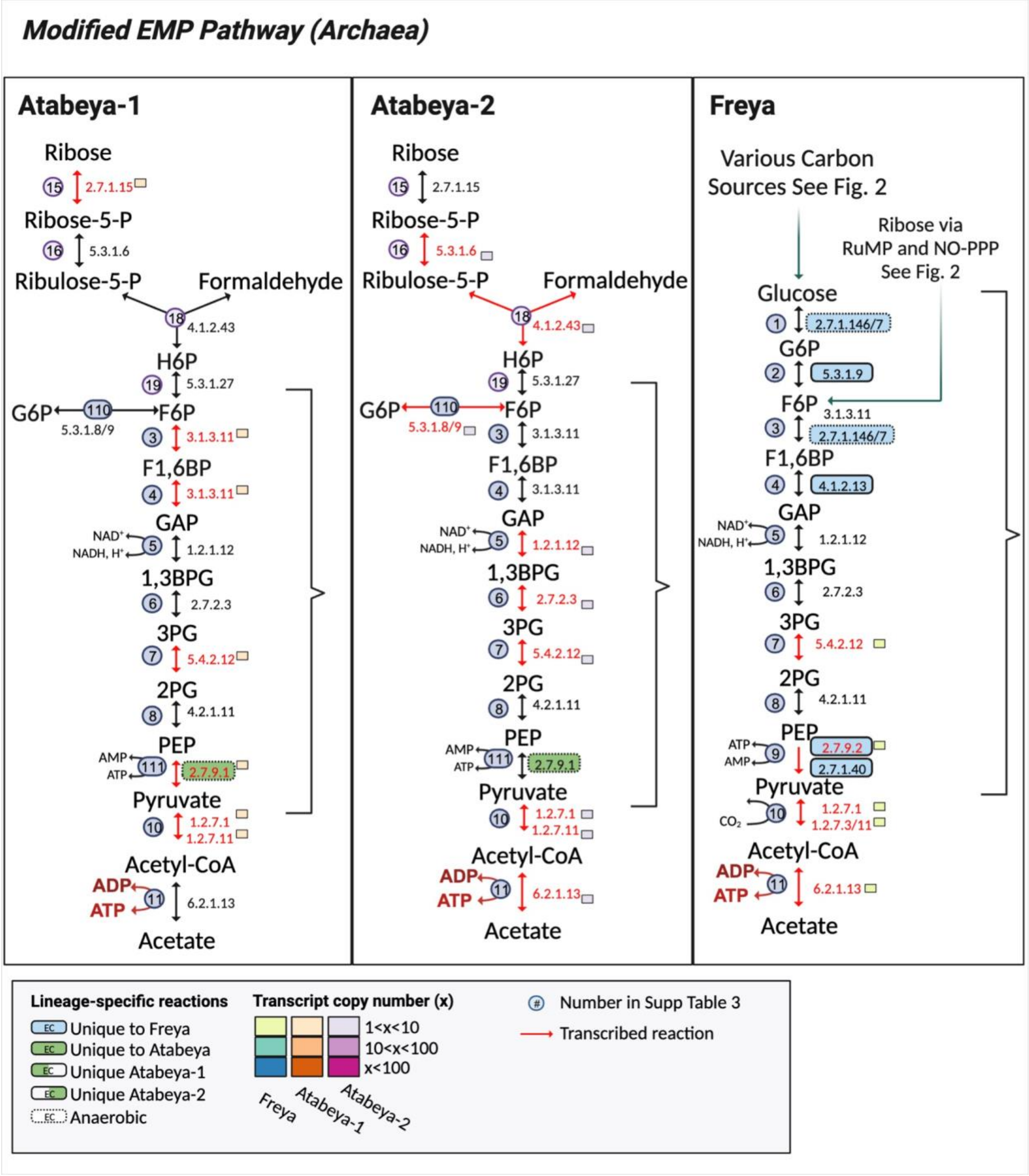

**Fig. S14:** Transcriptome map of the Ribulose Monophosphate Pathway (RuMP) and the Nonoxidative Pentose Phosphate Pathway (NO-PPP) for Atabeyarchaeia (Atabeya-1 and Atabeya-2) and Freyarchaeia (Freya). The numbers correspond to those in Figure 2 and Supplementary Table 7. Red arrows indicate mapped transcripts and the rectangles show the number of transcripts, matching Figure 2. Created using BioRender.com.

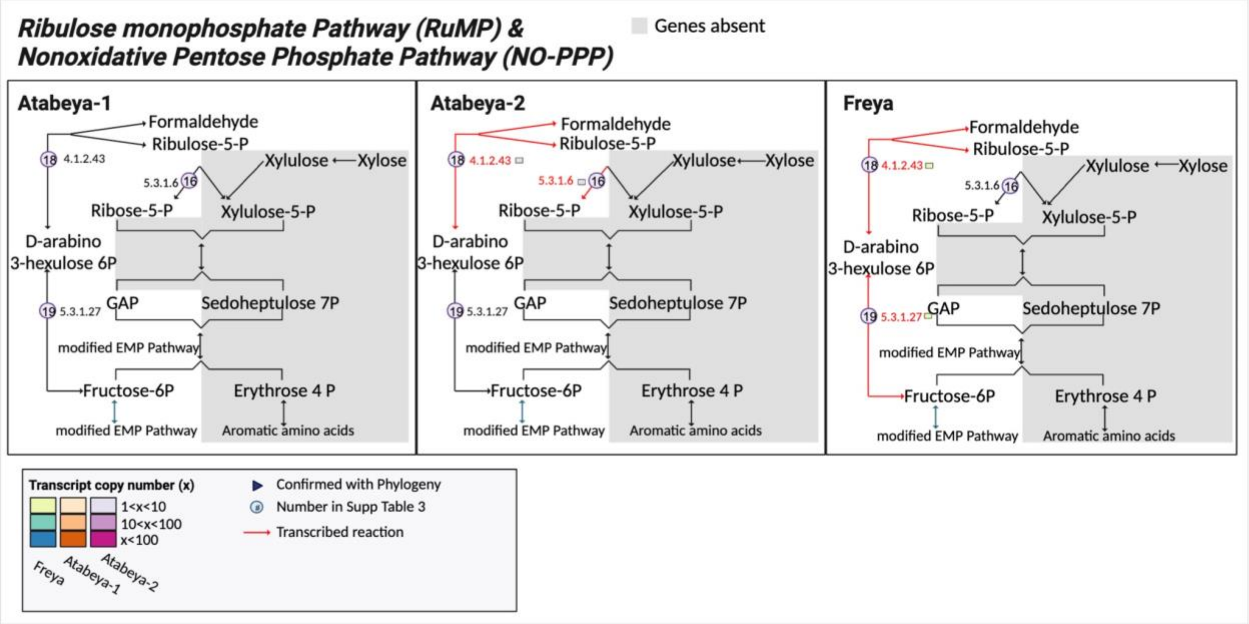

**Fig. S15** Maximum likelihood phylogeny of aldehyde oxidoreductases (AOR) to determine the specific family and potential substrate-specificity of the highlighted Atabeya-1, Atabeya-2, Freya sequences. The sequences for the three complete genomes were extracted and aligned with identified sequences from Arndt 2019. Mafft auto (v7.505) and trimAl -gt 0.5 (v1.4.rev15) were used for aligning and trimming, respectively. We used Iqtree (v1.6.12) to determine the maximum likelihood phylogeny, and LG+F+R4 was the model chosen according to BIC. Based on our analysis Freya and Atabeya sequences cluster with three known families (WOR5, XOR, and FOR) and two undescribed groups. Both Atabeyarchaeia genomes cluster with FOR and WOR5; whereas, Freya has sequences clustering with those groups and XOR. The family classification of Atabeyarchaeia and Freyarchaiea AORs is supported by the metabolism shown in **Figure 2**.

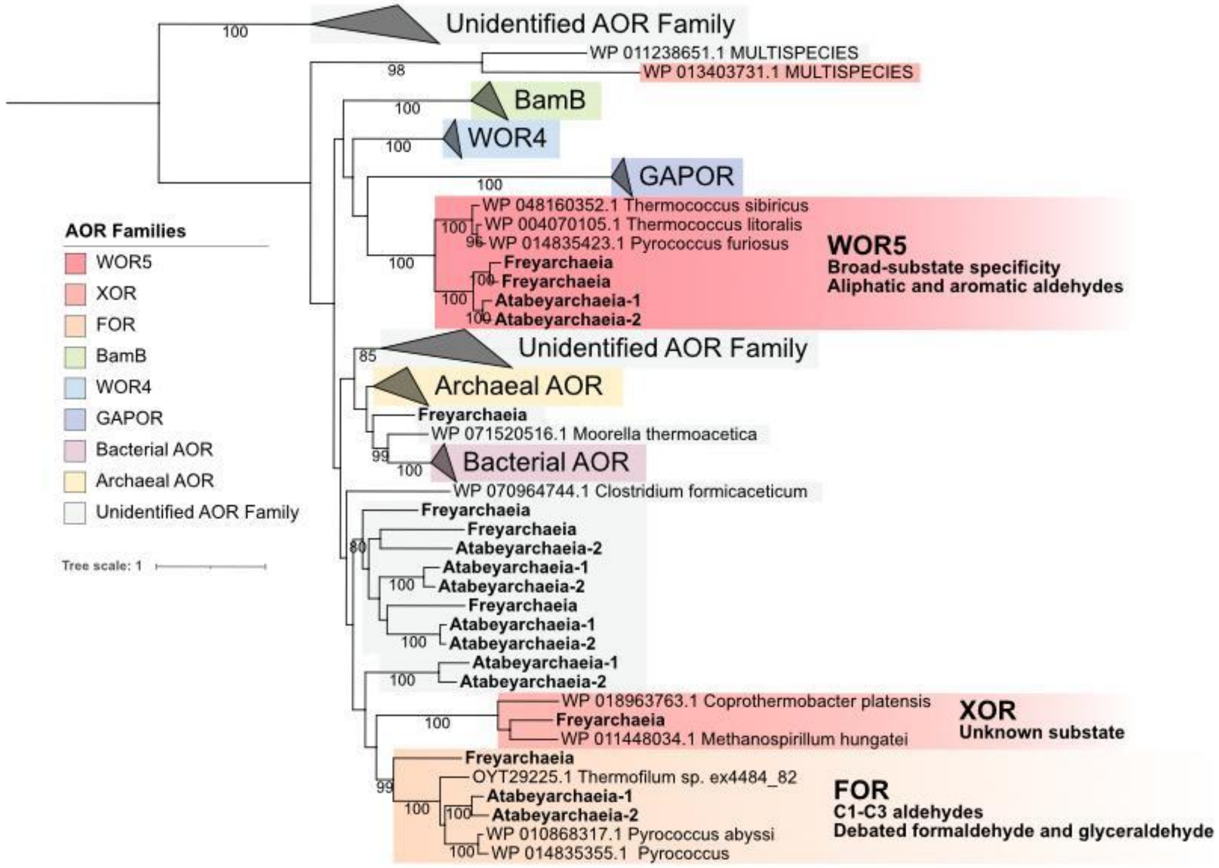

**Fig. S16:** Maximum likelihood phylogeny of RubisCo-like proteins (group IV and methanogenic group III). We identified two Atabeyarchaeia and two Freyarchaeia RbcL-like in the complete genomes, which were aligned with a subset of 1,000 references.. The sequences were aligned and trimmed with MAFFT v7.505 and trimAl v1.4.rev15. The tree was generated with IQ-TREE v.1.6.12, model LG+C20+R+F. Ultrafast bootstrap support values of >80 are shown.

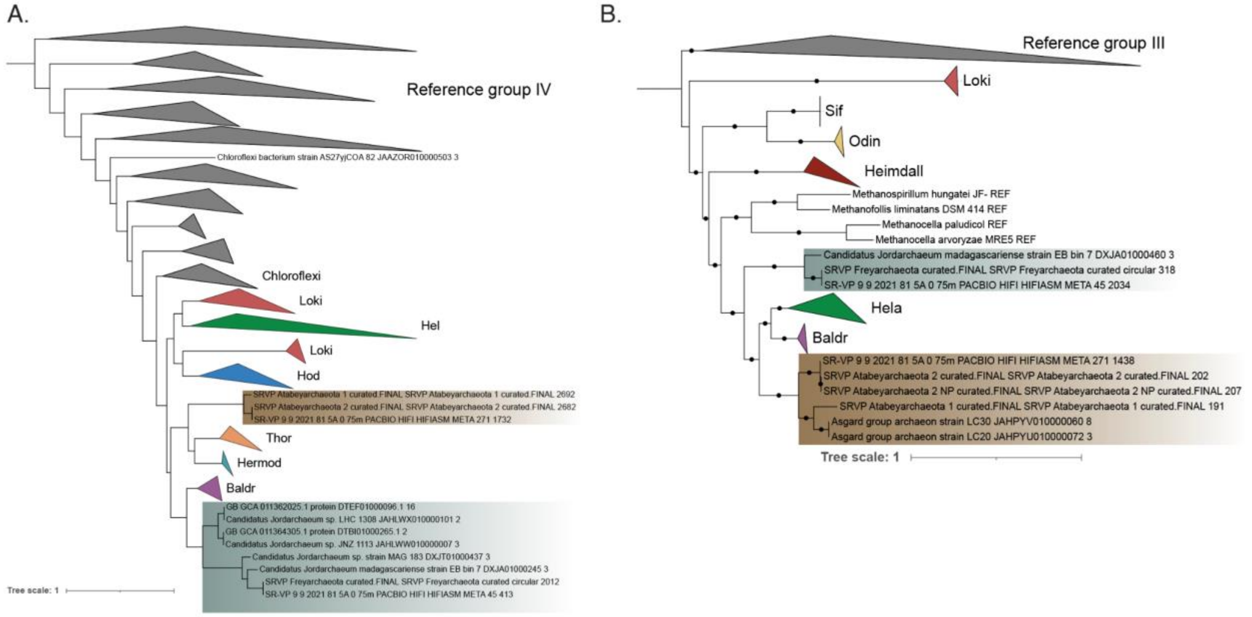

**Fig. S17:** Maximum likelihood phylogeny of GlpA. We identified two Atabeyarchaeia and two Freyarchaeia GlpA proteins in the complete genomes, which were aligned with a subset of 123 GlpA references from uniprot and 50 top hits from NCBI against the non redundant database in November 2023. The sequences were aligned and trimmed with MAFFT v7.505 and trimAl v1.4.rev15. The tree was generated with IQ-TREE v.1.6.12, model LG+C20+R+F. Ultrafast bootstrap support values of >80 are shown.

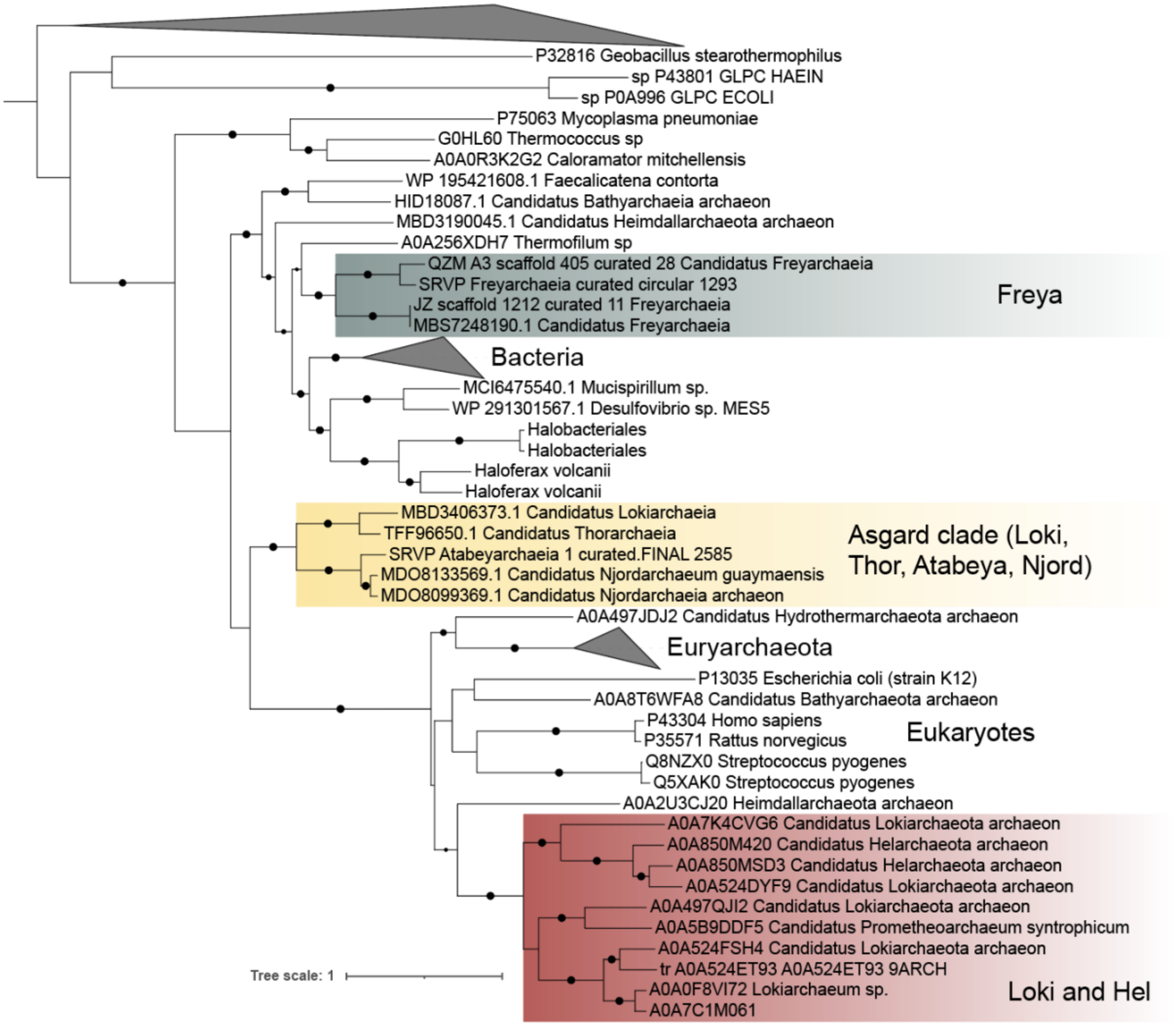

**Fig. S18**: Maximum likelihood phylogeny of catalases to determine the specific clade of the Atabeyarchaeia-1 and Atabeyarchaeia-2 sequences. The Freyarchaeia catalase was removed after several iterations from the tree as it was a partial sequence, grouping with Atabeyarchaeia-1 and Atabeyarchaeia-2 sequences. Initially, catalase clades were determined by subsetting the bacterial typical catalase sequences in https://doi.org/10.3389/fmicb.2021.645477 by group with MMSeq2 and aligning the subset with Mafft auto (v7.505). I augmented this alignment with reviewed sequences from IPR024711 (clade 1&3) and IPR024712 (clade 2). This expanded set of sequences was aligned with Mafft auto (v7.505) and trimmed with trimAl -gt 0.5 (v1.4.rev15). After several rounds of manually checking the alignment, we used Iqtree (v1.6.12) to produce the maximum likelihood phylogeny, and LG+R8 was the model chosen according to BIC. The clade classification of Atabeyarchaeia and Freyarchaeia catalase hits supported the annotation**s** in **table S7**. Ultrafast bootstrap support values of >80 are shown.

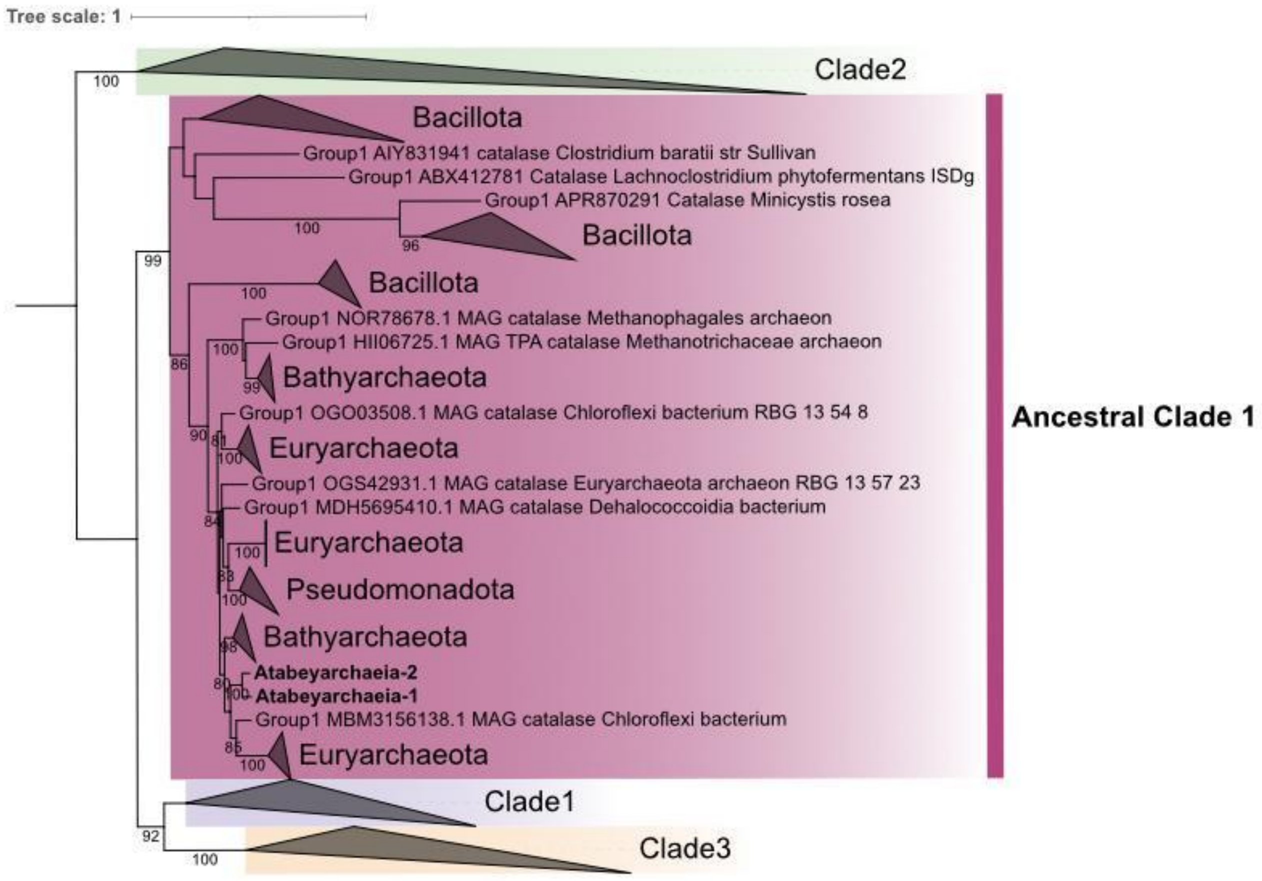

**Fig. S19**: Maximum likelihood phylogeny to check annotation of catalase peroxidase in Freyarchaeia. The highlighted Freya sequence was blasted against those peroxidase sequences in the PeroxiBase Database in October 2023. These sequences were downloaded, aligned with Mafft auto (v7.505), and trimmed with trimAl -gt 0.5 (v1.4.rev15). Iqtree (v1.6.12) was used to produce the maximum likelihood phylogeny, and WAG+I+G4 was the model chosen according to BIC. The clade classification of the Freyarchaeia catalase peroxidase hit was supported by the annotations in Table S7. Ultrafast bootstrap support values of >80 are shown.

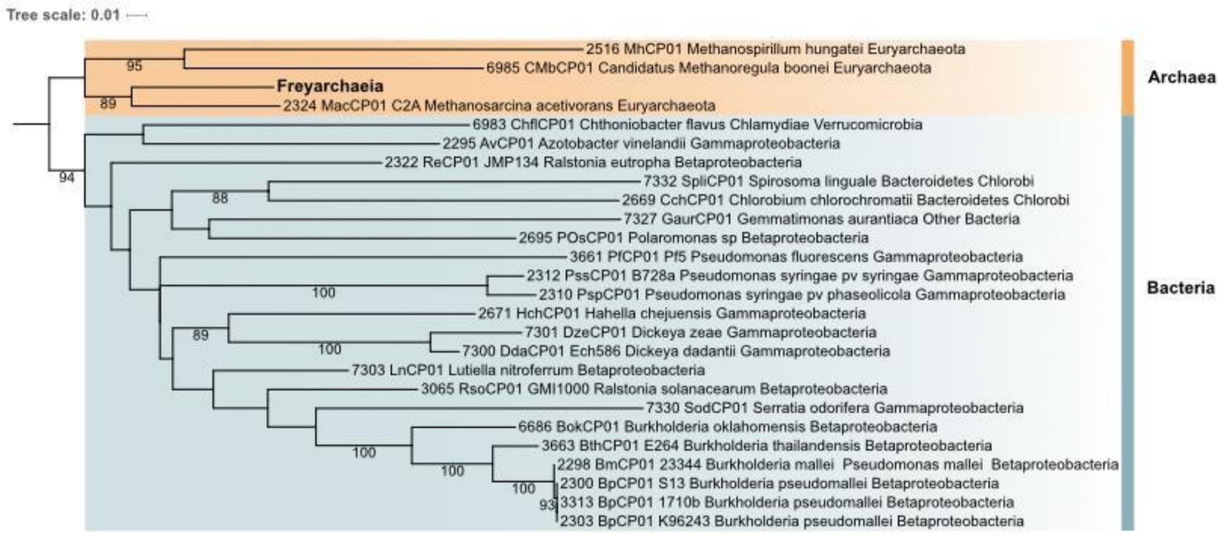

**Fig. S20**: Transcriptome map of the reactive oxygen species repair/damage pathways and hydrogen peroxide metabolic pathways for Atabeyarchaeia (Atabeya-1 and Atabeya-2) and Freyarchaeia (Freya). The numbers correspond to those in Figure 2 and table S7. Red arrows indicate mapped transcripts,the rectangles show the number of transcripts, and the green boxes behind the EC numbers indicate lineage-specific reactions for Atabeya genomes, matching **Figure 2**. Created using BioRender.com.

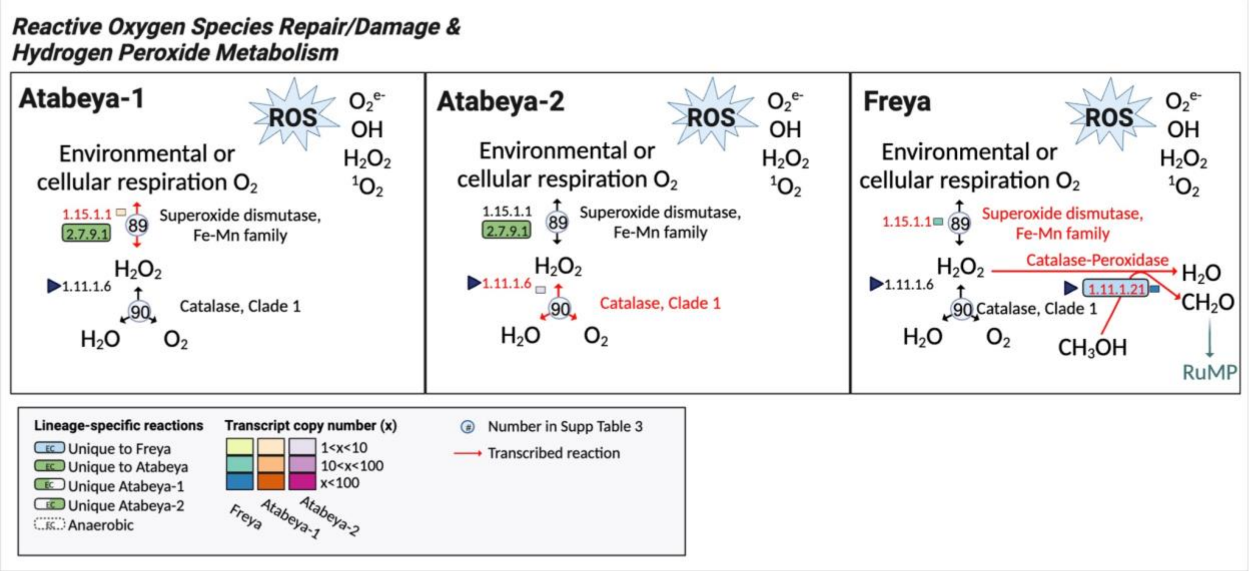

**Fig. S21** Phylogenetic analysis shows that the Sec elongation factor sequences from Atabeyarchaeia and Freyarchaeia are closely related to other Asgard members and Eukaryotes Ultrafast bootstrap support values of >90 are shown.

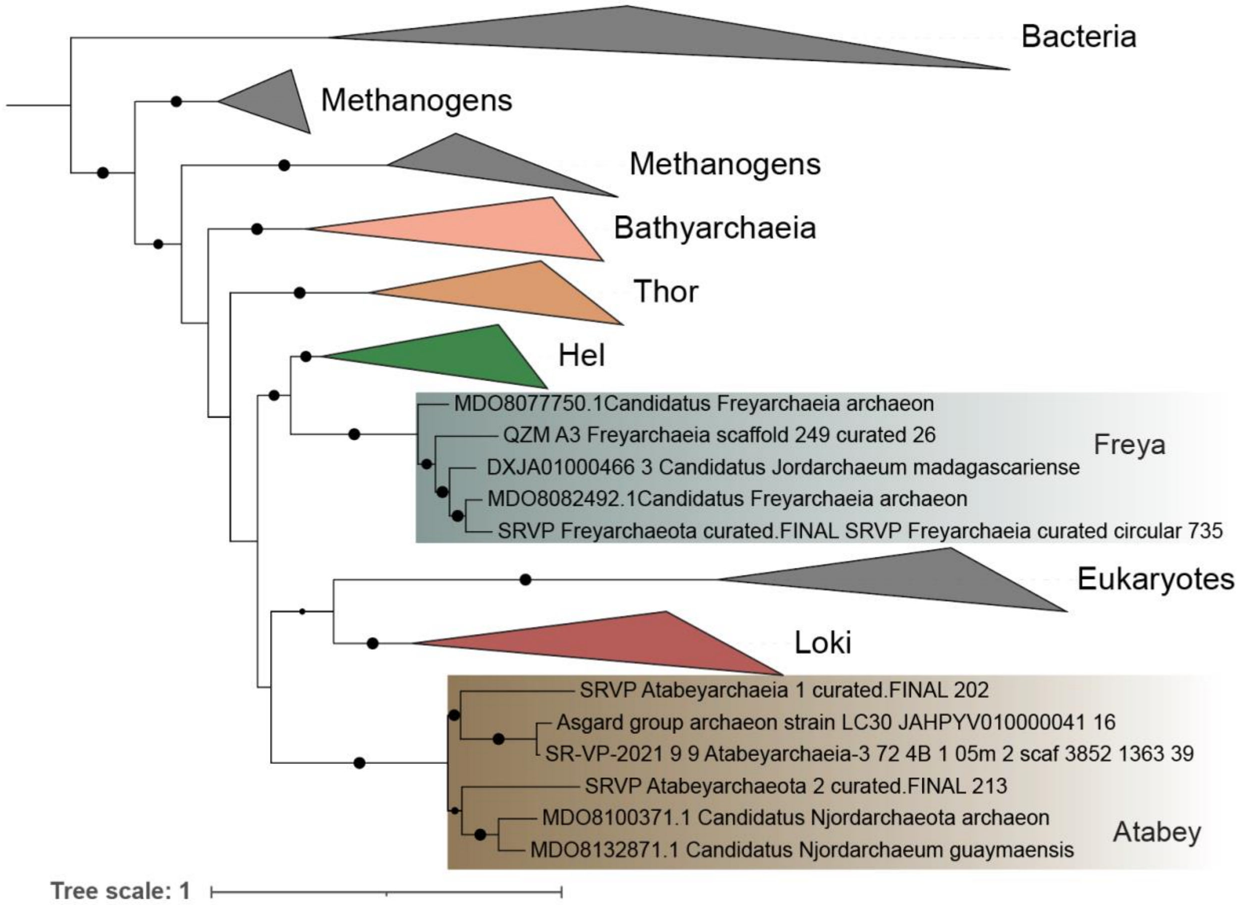

**Fig. S22** Maximum likelihood phylogeny of iron-containing alcohol dehydrogenases (ADH) in Atabeyarchaeia and Freyarchaeia complete genomes to determine potential substrate specificity of these enzymes. Freyarchaeia has a putative NADPH-dependent butanol dehydrogenase (BDH) similar to that in Chloroflexi bacterium (PIW40290), which is shown in Figure 2 and discussed in detail in the "**Extended Metabolism**" section. Ultrafast bootstrap support values of >80 are shown.

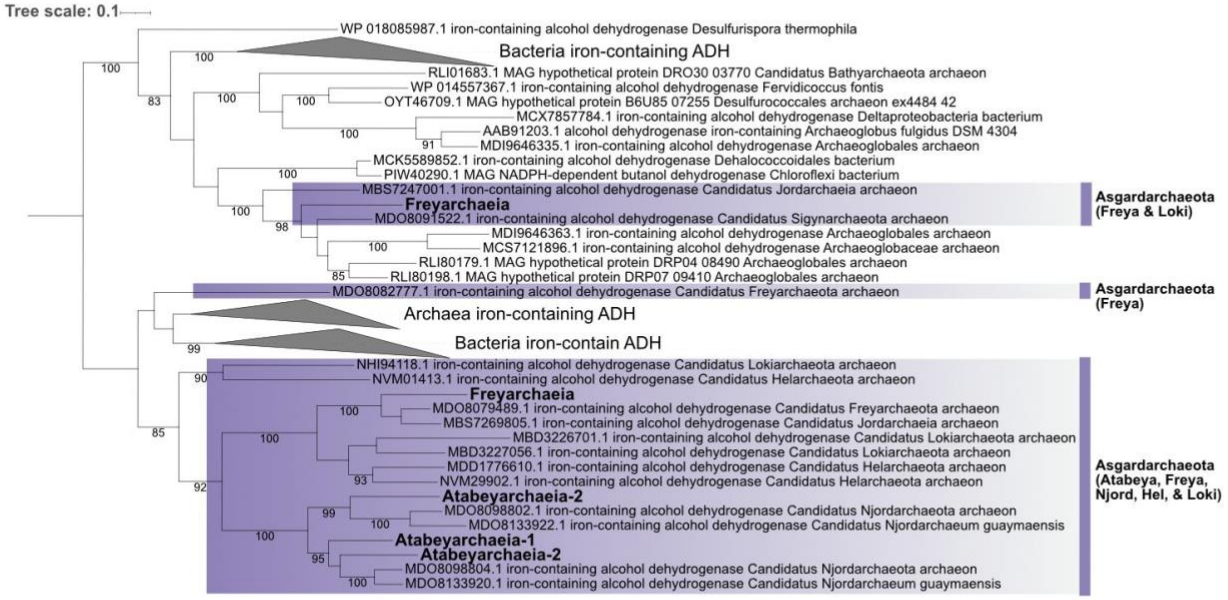

**Fig. S23:** Transcriptome map of the proposed butanol oxidation pathway for Freyarchaeia. The numbers correspond to those in Figure 2 and Supplementary Table 7. Red arrows indicate mapped transcripts and the rectangles show the number of transcripts, matching Figure 2. Created using BioRender.com.

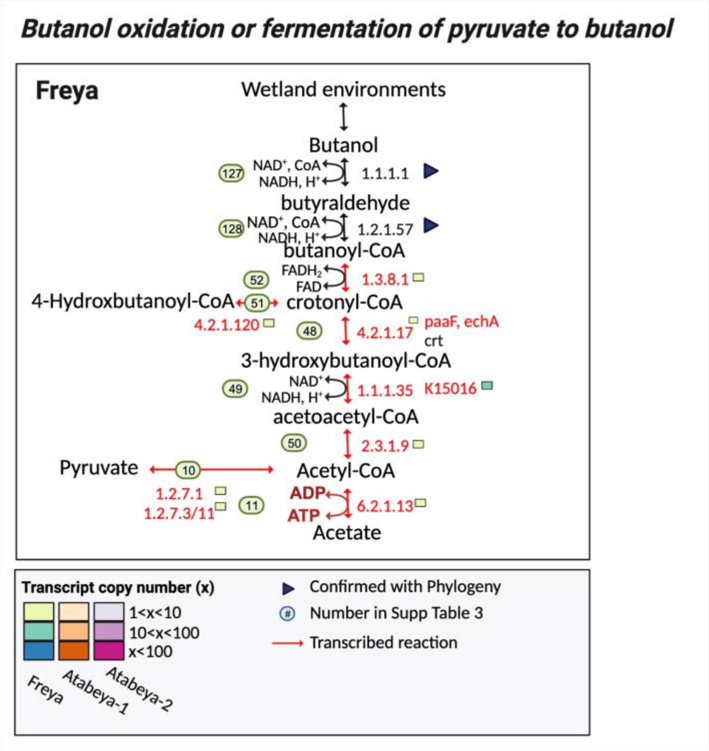

**Fig. S24** Maximum-likelihood tree, inferred with IQtree and the best-fit LG+F+R10 model, using Phylosift 37 markers from Asgardarchaeota and TACK. Ultrafast bootstrap support values of >80 are shown.

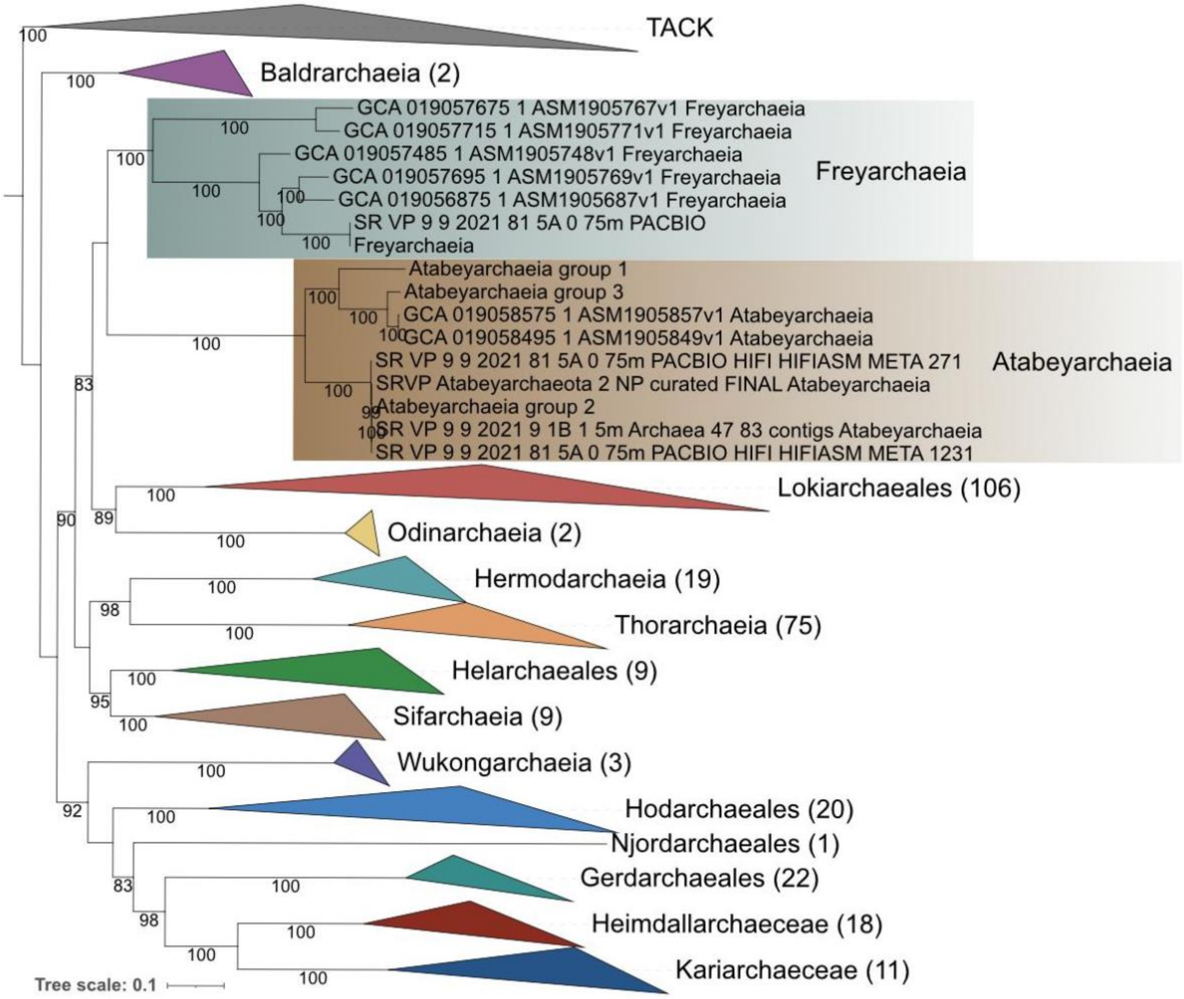

Supplementary Table 1 Description: *Excel File* Overall genomic statistics of complete genomes, MAGs and 245 public Asgard reference genomes downloaded from Bacterial and Viral Bioinformatics Resource Center (BV-BRC), database on March 20th, 2022 BCR

Supplementary Table 2 Description: *Excel File* Average amino acid identity (AAI) comparision of Atabeyarchaeia and Freyaerchaeia genomes and phylogenetically related Asgardarchaeota phyla.

Supplementary Table 3 GTDB version 2.3.0: These results shows our genomes but also publicly available genomes used for the comparative genomic analyses

Supplementary Table 4 Description: *Excel File,* The tRNA genes and introns of Atabeyarchaeia and Freyaerchaeia genomes.

Supplementary Table 5 Description: *Excel File*. Eukaryotic signature proteins in Atabeyarchaeia and Freyarchaeia

Supplementary Table 6 Description: *Excel File*, Metatranscriptomes

Supplementary Table 7 Description: *Excel File*, Metabolism sheet: Cell Diagram sheet includes key metabolic genes identified in Atabeya and Freya complete genomes shown in Figure 2 metabolic overview. The sheet includes information about number of identified gene copies, metatranscript copy number, psortv 3.0.3 locations, and if there is a corresponding supplementary phylogeny. The next nine sheets contain raw annotations for KofamKOALAv1.3.0, Interproscanv5.6.1-93.0-64, MEBsv2_Pfamv34.0, HADEG_Alkanes, METABOLICv4, dbcan, MEROPSv12.4, DRAM, and HyDB, respectively

Supplementary Table 8 Description: *Excel File*, Hydrogenases identified in Atabeyarchaeia and Freyarchaeia complete genomes.

Supplementary Table 9 Description: *Excel File,* PSI-Blast results from ArCOGS

Supplementary Table 10 Description: *Excel File,* The identity and coordinates of the selenocysteine machinery in the Atabeyarchaeia and Freyaerchaeia genomes.

Supplementary Table 11 Description: *Excel File,* The identity and coordinates of the selenoproteins and their insertion sequences in the Atabeyarchaeia and Freyaerchaeia genomes.
